# Maternal exposure to prostaglandin e2 results in abnormal dendritic morphology in the cerebellum and related motor behaviour in mouse offspring

**DOI:** 10.1101/2023.10.12.562077

**Authors:** Ashby Kissoondoyal, Kelly Ho, Christine Wong, Dorota A. Crawford

## Abstract

The lipid signalling molecule prostaglandin E2 (PGE2) is important in healthy brain development. Abnormal PGE2 levels during prenatal development, which can be influenced by genetic causes and exposure to various environmental risk factors, have been linked to increased prevalence of Autism Spectrum Disorders (ASDs). Growing research in animal models aims to provide evidence for the mechanisms by which increased or reduced PGE2 levels influence brain development. In this study, we show that maternal exposure to PGE2 in mice at gestational day 11 (G11) results in molecular changes within the cerebellum and associated behaviours in offspring. We observed a decrease in cerebellar cell density originating at G11 (in males and females) and at G16 (in females only). In Golgi-COX-stained cerebellar slices from PGE2-exposed offspring at the postnatal day 30 (PN30), we found an increase in dendritic arborization, the odds of observing dendritic loops, dendritic spine density, and the odds of observing mature (mushroom-shaped) spines. We also observed a decrease in the expression level of the cytoskeletal protein β-actin, the actin associated protein spinophilin, and the cell adhesion protein N-Cadherin. In addition, we found that specifically PGE2-exposed male offspring exhibited abnormal cerebellar related motor function. This study adds further evidence that changes in the PGE2 levels during critical times may impact the developing brain differently in males and females. These findings also emphasize the importance of examining sex differences in research relevant to neurodevelopmental disorders.

## Introduction

Prostaglandin E2 (PGE2) is a bioactive lipid signalling molecule, that plays a vital role in the early nervous system development, including mediating cell proliferation and differentiation, dendritic spine formation, synaptogenesis, and learning and memory [1, 2]. Synthesis of PGE2 begins with the release of arachidonic acid (AA) from the cell membrane through the enzymatic activity of phospholipase A2 (PLA2) [3]. PGE2 is then metabolized from AA by cyclooxygenase -1 or -2 (COX-1 or COX-2) [4, 5]. COX-1 and COX-2 are constitutively expressed in the brain with COX-1 primarily within microglial cells [6, 7] and COX-2 mainly in neuronal cells [8, 9].

We have previously reviewed the existing evidence showing the link between abnormal COX-2/PGE2 signalling and the etiology of Autism Spectrum Disorders (ASDs) [10–12] PGE2 levels can be affected during prenatal development by genetics [13], or exposure to various environmental risk factors, including air pollution [14, 15], glyphosates and other herbicides [16, 17], phthalate molecules used in consumer products [18–20], and common over the counter drugs such as acetylsalicylic acid and acetaminophen [21–29], all shown to be associated with ASDs in humans. Misuse of the drug misoprostol, a synthetic analogue to PGE2, for a termination of pregnancy also leads to ASD and Moebius syndrome [30, 31]. These exogenous chemicals can diffuse or be shuttled through the blood brain barrier and the placenta during a critical time in pregnancy and impact the PGE2 levels in the developing brain [32]. Despite the clinical and epidemiological evidence showing that the COX2/PGE2 signalling pathway has etiological significance to the pathogenesis of ASDs, the molecular mechanisms by which changes in prenatal PGE2 levels influence brain development are still not well understood largely due to the lack of proper experimental model systems. *In vitro* studies in our lab provided first evidence in neuronal cells for the connection between PGE2 signalling and neuronal morphology. For example, an elevated level of PGE2 in various neuronal cell lines increased calcium levels in the cytosol and growth cones, and affected neural stem cell proliferation, migration and differentiation [33–36]. In our *in vivo* models – mice offspring prenatally exposed to PGE2 (PGE2-exposed) and mice lacking COX-2 (COX-2^-^-KI) – exhibit autism-related behaviours such as deficits in social communication, repetitive/restrictive behaviours, and anxiety in a sex-dependent manner [37–39]. We also found a differential expression of developmental genes involved in biological processes such as dendritic pathfinding and synaptic function [38, 40].

ASDs include a group of complex neurodevelopmental disorders, which are primarily characterized through behavioural deficits [41] with males 4 times more likely to be diagnosed than females [42]. Beside the core symptoms ASD individuals display motor deficits, including delays in developmental milestones including lying, righting, sitting and crawling [43], deficits in both fine and gross motor movements [44], difficulties in postural control [44, 45], and a general lack of motor coordination appearing as clumsiness [46–48]. These atypical behaviours often correspond to molecular and morphological changes within the cerebellum [49]. Early pre- and postnatal disruptions of cerebellar circuitry, either through injury during birth or developmental abnormalities in the development of the cerebellum are correlated with later diagnosis of ASD [50–53]. Other studies in experimental models have reported that an early postnatal increase or decrease in PGE2 levels results in abnormal cerebellar development and decreased reciprocal social behaviour in male rats [54, 55]. Our previous study in COX-2^-^-KI mice also showed sex-dependent changes in the morphology of dendrites and dendritic spines within the cerebellum [56].

In this study we examined the effect of maternal exposure to PGE2 (in PGE2-exposed mice) at gestational day 11 (G11) on sex-dependent morphological changes within the cerebellum and associated behaviour in offspring. We first quantified density of cells originating from G11 and G16 critical timepoints (the onset and late neurogenesis stages) as well as dendrite length, arborization, and looping as well as dendritic spine density and shape at postnatal day 30 (PN30). We also quantified the expression levels of β-actin (cytoskeletal protein), spinophilin (actin-binding protein enriched in dendritic spines), and N-cadherin (synaptic cell adhesion protein). Finally, we evaluated whether the molecular changes observed can be coupled with deficits in cerebellum related motor coordination by implementing the adhesive sticker, grid walking and cylinder tests. We show for the first time *in vivo* that a single exposure to PGE2 during a critical time in pregnancy has significant molecular and morphological consequences within the cerebellum, that result in sex-dependent manifestations of relevant behaviours in offspring.

## Methods

### Animals

Male and female C57BL/6 mice were purchased from Jackson Laboratories. All animals were housed via group housing at the York University animal facility. Male and female C57BL/6 mice were housed together overnight for breeding and each morning females were checked for the presence of a vaginal plug. Gestation day 1 (G1) was marked as the day that a vaginal plug was observed and from that point on the females were housed individually for the remainder of their pregnancy. On G11, females were injected with a single subcutaneous injection of 0.25µg/g 16,16 dimethyl prostaglandin E2 (dmPGE2; Cayman Chemical) diluted in saline to a final volume of 300µL. While considered an analogue to PGE2, dmPGE2 is metabolized at a slower rate than PGE2 and as such remains active for a longer time frame [57, 58]. Maternal PGE2 can access the developing brain through diffusion or specific transporters across the blood brain barrier and the placenta [59–61]. Control animals were injected with 300µL saline. Injections of dmPGE2 were performed on G11 to coincide with the onset of neurogenesis in mice [62]. In the mouse, the neurogenic interval extends from gestation day 11 (G11) through early G17 in most brain regions [63–66]. It also corresponds to the time the drug misoprostol was misused by pregnant women to terminate pregnancy that resulted in Moebius syndrome and autism [30, 67]. Mouse offspring subjected to dmPGE2 exposure are referred to as “PGE2-exposed mice” and those injected with saline as “Saline control mice”.

### CldU and IdU labelling

Subcutaneous injections of 5-Chloro-2’-deoxyuridine (CldU) or 5-Iodo-2’deoxyurdine (IdU) (Sigma) were administered (50μg/g dissolved in saline) to pregnant saline control and PGE2-exposed mice. CldU and IdU are different thymidine analogues that are incorporated into the dual helix of any cell actively synthesizing DNA during at the time that the thymidine marker is injected [67, 68] With this technique, we were able to detect differences in cell proliferation in the cerebellum at two unique time-points. An injection of CldU at G11 and IdU at G16 were given, to capture both early and late phases of this interval, and animals were sacrificed at Postnatal Day 8 (PN8). For PGE2-exposed mice, pregnant dames were given a single co-injection of CldU with PGE2 at G11 at described concentrations.

### Immunohistochemistry

Left hemisphere brain samples were extracted and immediately fixed at 4°C for 48 hrs in 4% paraformaldehyde (in PBS). Paraffin-embedding and serial slicing from the mid-sagittal plane outwards in slices of 4μm in thickness were completed by The Centre for Phenogenomics (Toronto, Canada). Immunohistochemistry was performed as previously described [69]. Xylene incubation was performed to remove paraffin from samples. This was followed by rehydration through serial ethanol incubation and followed by cell permeabilization with 0.2% Triton X-100 in PBS. We then performed antigen retrieval using 0.01M pH 6.0 sodium citrate buffer followed by a 1.5N HCl incubation. Sections were circled using a liquid blocking super PAP pen (Cedarlane) and sections were then treated at 37°C for 3 min with 0.25% trypsin EDTA in a pre-warmed hydration chamber. The remaining steps were then performed in a hydration chamber. For primary antibody staining, samples were first blocked in 5% goat serum diluted in PBS, followed by 4°C overnight incubation of the Cldu specific antibody (Rat anti-BrdU; 1:100, ab6326, Abcam), diluted in 5% goat serum in PBS. Samples were then incubated at 37°C with agitation for 20 min at 225 rpm in a high stringency wash of low salt TBST buffer (36 mMTris, 50mM NaCl, 0.5% tween-20; pH 8.0) [68]. For the Idu primary antibody (Mouse anti-IdU;1:100, ab181664, Abcam), samples were incubated overnight at 4°C overnight. Secondary antibody incubation was completed in the dark with Alexa Fluor 555 goat anti-mouse (1:500, ab150118, Abcam) and Alexa Fluor 488 goat anti-rat goat anti-rat (1:500, ab150165, Abcam). Coverslips were mounted with ProLong Gold Antifade Mountant (ThermoFisher). CldU and IdU staining was visualized and captured using an Eclipse 80i upright fluorescent microscope with DS-5MC camera (Nikon). To correct for variations between brain slices, estimated cell density measurements in the cerebellum were calculated by dividing the total number of cells counted by the area.

### Golgi-COX staining

Whole brain tissues were collected at PN30 from 3 males and 3 females Saline Control and PGE2-exposed mice offspring from 3 separate litters from both. Before staining, the cerebellar hemispheres were separated in half using the central vermis as a reference. Golgi staining was performed as we have previously described [70]. After extraction, brains were dissected in half, rinsed in cold 0.1M PBS solution and immediately placed in a 4% paraformaldehyde solution for 24h. Following fixation brain samples were washed with ddH2O and placed in Golgi-COX solution (5% w/v potassium dichromate, 5% w/v mercuric chloride, and 5% w/v potassium chromate) dissolved in ddH2O. After 24h the brain tissues were transferred into fresh Golgi-COX solution and incubated for an additional 7 days at room temperature in the dark. The brains were then transferred into tissue protectant solution (30% w/v sucrose, 20% w/v ethylene glycol, and 1% w/v polyvinylpyrrolidone) in ddH2O and kept in this solution for up to 7 days.

For Golgi-COX staining brains were sliced along the sagittal in slices of 100µm in thickness using cryostat sectioning by the research histology lab at the University Health network (Toronto, Ontario; www.uhnresearch.ca). Brains were sectioned from the median plane by the same technologist. Trimming laterally from the central vermis was performed prior to sectioning and at least 3 serial sections were collected from each animal for analysis. Brain slices were mounted onto gelatin-coated slides and staining was developed with a 3:1 ammonia to H2O solution followed by a 5% w/v sodium thiosulfate in H2O solution. Serial dehydration in 70%, 95% and 100% ethanol and then immersion in xylene was performed before the slides were cover-slipped.

Golgi-COX-stained slides were imaged on the Zeiss Laser Scanning Confocal Microscope (LMS 700; Advanced Light and Electron Microscopy at York University), using brightfield microscopy. The full cerebellum of each animal was imaged at 10x magnification with z-stacks taken every 1-2.5µm (50-100 stacks on average). Individual cell within the cerebellum were imaged using 100x magnification with z-stacks taken every 0.01 – 0.02µm (1000-2000 stacks on average). Cells were selected from the posterior lobe of the cerebellum within the intermediate zone, with attention made to ensure an even distribution of cells were chosen across each slide.

Analysis of cell density and dendritic arborization, length, and loop quantification

Cerebellar G11 and G16 cells were counted, and density was calculated by dividing the total number of CldU and IdU cells by the area of interest as described in [38, 39]. Image analysis was completed using NIS-Elements software (Nikon). Cell selection, sholl analysis and quantification of dendritic loop formation was performed as previously described in [56]. From the 10x magnification images taken, 5 neuronal cells per animal within the cerebellum were chosen for a total of N=60 cells with 15 cells per condition. Cells were selected from the posterior lobe of the cerebellum. The Simple Neurite Tracer (SNT) plugin within the open-source Fiji ImageJ software was used to conduct sholl analysis [71–73]. Concentric circles were generated by the software every 10µm from a point defined in the center of the soma of each neuron examined and the number of intersections cell extensions made with these circles was measured. *Dendritic length* measurements were taken from SNT established arbors. The average length of all primary dendrites was taken for each cell used for sholl analysis. We previously demonstrated a method for the quantification of self-fasciculation in dendrites in the cerebellum of COX-2^-^-KI mice [74]. Using the traced dendrites from the SNT plugin, the maximum angle of each dendrite was used to classify each dendrite as looping or non-looping; dendrites with a turning angle greater than 270° were classified as looping as previously described [56].

### Dendritic spine analysis

Individual cell images were used to quantify dendritic spine density and morphology. 10 cells were selected from each of 3 mice for a total of 30 cells per condition. Dendritic spines were measured as any extension coming from a dendrite. Identification and measurement of dendritic spines was performed using Nikon NsAi software (Nikon). Training of the SegmentAI software was performed on a previously quantified image set with a training loss of less than 0.02 used as a cut-off for successful training. DenoiseAI was used to reduce background detection. After SegmentAI detection of dendritic spines, false positives were filtered out by length (> 0.2µm), width (> 0.2µm), circularity (< 0.88), and area (> 0.6µm^2^). Dendritic spine density was calculated as the number of AI identified dendritic spines per the length of the dendrite (which was determined semi-automatically for each dendrite) [75, 76]. Each dendritic spine was classified as a shape through a sequential top-down approach we have previously described [74]. Spines with a width > length were classified as stubby, of the remainder those with a width > 0.6µm were classified as mushroom, and those that remained were classified as thin.

### Motor behaviour

Mice were kept on a 12-hour light/dark cycle with behavioural testing administered from the start of the light phase. Mice were separated by condition into PGE2-exposed (see maternal injections in methods) and vehicle (Saline), and by sex (male and female) on the day of weaning. From each condition 10 males and 10 females were selected from at least 3 litters. Between behavioural testing, all equipment was disinfected and deodorized with antiseptic clinicide and wiped clean. All behavioural tests were administered by the same female researcher to avoid increases in stress levels reported in behavioural tests administered by male experimenters [77]. To facilitate more accurate comparisons of overall motor behaviour of each mouse, each mouse was tested in each of the adhesive sticker, grid walking and cylinder tests. All behavioural tests were measured on PN30 with training days occurring the two days prior (PN28 and PN29) and recorded. The recorded videos were analyzed by 5 trained individuals.

#### Adhesive sticker test

The adhesive sticker test was used to assess sensory motor limb coordination [78–80]. A red sticker with a diameter of 0.5cm was placed just above the nose of a restrained mouse. The mouse was released into an empty cage with bedding and the mouse was recorded from above. Each mouse was tested at PN30 for 3 trials with at least 2 minutes of rest between each trial. The average of their measurements for the *first contact* with the adhesive sticker, and the number of *swipes per second* to remove the sticker were recorded. Time until first contact was recorded as the time from when each mouse was released until either forelimb made contact with the adhesive sticker, and number of swipes per second were calculated as the number of attempts at removing each sticker divided by the time from *first contact* until the sticker was removed.

#### Grid walking test

The grid walking test was used to assess sensory motor coordination of the limbs and balance [81, 82]. We used an apparatus consisting of an elevated metal grid, enclosed with wooden walls and an open top. An opening in the mesh grid on one end allowed the mouse to escape into a home cage with bedding and space to hide. The grid walking test spanned 3 days (2 training and one testing) starting on PN28 (testing day occurred on PN30). Training days consisted of two sets of 5 runs in which the mice were placed inside the grid walking apparatus (60cm x 13cm x 7.5cm) with a mesh grid of 1cm x 1cm to acclimatize to the test. Testing day included 1 warmup run followed by three trials with at least 2 minutes of rest between each trial. The testing apparatus (60cm x 20cm x 7cm) had a mesh grid of 2.5cm x 2.5cm. To record limb slips, a mirror was placed such that the bottom of the mesh grid could be recorded. The recorded videos were manually analyzed by 5 trained individuals and the average of their measurements for number of slips through the grid and the total time to escape were recorded.

#### Cylinder test

Exploratory behaviour and spatial motor behaviour were assessed using the cylinder test [83–85]. Mice were placed into a long glass cylinder (8.5cm diameter x 21.5 cm) with an open top. The glass cylinder was placed on top of a pane of glass and the bottom view was recorded. The test spanned 3 days (2 training and 1 testing) starting on PN28 (with testing occurring on PN30). Training days consisted of 3, 90 second runs with at least 5 minutes between each run. On testing days, each mouse performed 1 warmup run of 3 minutes, followed by 3, 10-minute trials, with at least 5 minutes rest between each trial. Recorded videos were manually analyzed by 5 trained individuals and the average of their measurements for number of rears, steps, and forelimb glass touches were recorded.

### Protein isolation and western blotting

Cerebellar brain tissue samples were collected from Saline and PGE2-exposed male and female mice offspring from the same animals that underwent behavioral testing at PN30. Following homogenization of tissue samples using a polytron power homogenizer, total protein isolation was performed through standard RIPA method (Abcam). From each sample 30-40µg of protein was loaded into a 12% polyacrylamide gel as previously described [86, 87]. Protein samples from Saline Control and PGE2-exposed males and females were pooled for the analysis. In addition, protein samples from select PGE2-exposed male mice which performed exceptionally low or high on behavioral tests were loaded individually. Protein samples were separated using PAGE gel electrophoresis before being transferred to a 0.45µM nitrocellulose membrane (BioRad). Before probing with primary antibodies, membranes were washed in TBS and blocked in 5% milk in TBS-T (1X Tris buffer saline 0.05% Tween 20) for 1h at room temperature. Protein samples were probed with polyclonal rabbit anti-spinophilin (Abcam, 1:1000, ab18561, Cambridge, MA, USA) overnight at 4°C, and anti-β-Actin (Abcam, 1:5000, ab6276, Cambridge, MA, USA), anti-N-cadherin (Abcam; 1:10000, ab76011, Cambridge, MA, USA), and anti-GAPDH (Abcam; 1:5000, ab8245, Cambridge, MA, USA) for 1h at room temperature. After each primary antibody, membranes were washed 5 times in 1x TBS-T before probing with appropriate HRP-tagged secondary antibodies. Goat-anti-rabbit (Abcam, 1:10000, ab6276, Cambridge, MA, USA), and Goat-anti-mouse (1:10000, ab9789, Cambridge, MA, USA) were each incubated in 2% milk in TBS-T for 2h at RT. Probed membranes were then imaged with the Geliance 600 imaging system (Perkin Elmer). For quantification, protein signal intensity was first normalized to GAPDH signal intensity, and then relative protein expression was normalized to the expression of that protein in Saline control males (protein expression in Saline-M = 1).

### Statistics

All data collection and image and behavioural analyses were performed blind to the condition (PGE2-exposed vs Saline) and sex of the mice. Blinding was removed prior to statistical analyses. Statistical analyses were performed using the core open-source software R [88]. Categorical data (loop formation and dendritic spine shapes) were analyzed using multinomial logistic regression with all data presented as odd ratios (95% confidence intervals) in tables [89]. All remaining data including the remainder of dendritic morphology, all behavioral data, and all protein analysis were analyzed using linear mixed effect modeling to account for confounding variables in our data [90]. Variables of interest were assigned as main effects and confounds were assigned as random effects. Random effects in this study included litter, trial number, and technical replicates. While individual observations were taken from each neuron, each cell was assigned to a uniquely identified mouse (in which no mouse shared the same litter, condition, and sex qualifiers). During model determination, this accounted for repeated measurements within each mouse (biological replicate), while providing an appropriate power to our findings. Linear mixed models were fit by maximum likelihood and the Akaike Information criteria was used to determine the model of best fit for each data set. T-tests were performed with Satterwaite’s adjustment with significance determined as p < 0.05 for all tests. Data was displayed using a violin plot to depict the probability distribution of the data within a given treatment. All measurements were taken from a minimum of 3 independent biological replicates (ie litters) per treatment. The probability distribution represented in violin plot is based on measurements from the specified number of cells indicated above. Total sample sizes for each test were calculated based on literature using G*Power 3 software [91] using an estimated effect size of 0.25.

## Results

### Cerebellar cell density

We have previously demonstrated that maternal exposure of dmPGE2 at G11 altered expression of autism-linked genes and resulted in autism related behaviours in mice offspring [38, 39]. Here we observed that PGE2-exposed male and female offspring have a distinct localization pattern of CldU-labelled (G11) and IdU-labelled (G16) cells within the developing cerebellum at PN8. We separately counted cells originating from G11 and G16 (Fig 1A). A linear mixed effects model was used to fit G11 cell density (cells/mm^2^). Litter was assigned as a random control, and the fixed effects of condition and sex, as well as the interaction between condition and sex were examined. The interaction between condition and sex (t(7.3) = -2.558, p = 0.0363), condition (t(12.2) = -4.112, p = 0.0014), and sex (t(8.0) = 6.804, p = 0.0001) all had a significant effect on G11 cell density. Given the significance of the interaction term and the main effects we performed further pairwise comparisons. PGE2-exposed mice had a lower cell density than Saline control mice in males (Fig 1B, p = 0.0014, Saline =339.588, PGE2 = 156.256), and females (Fig 1B, p < 0.001, Saline = 595.015, PGE2 = 267.592). Within conditions, males had a lower cell density than females in Saline control mice (Fig 1B, p = 0.0001, M = 339.588, F = 595.015) as well as PGE2-exposed mice (Fig 1B, p = 0.0377, M = 156.256, F = 267.592).

**Figure 1.**
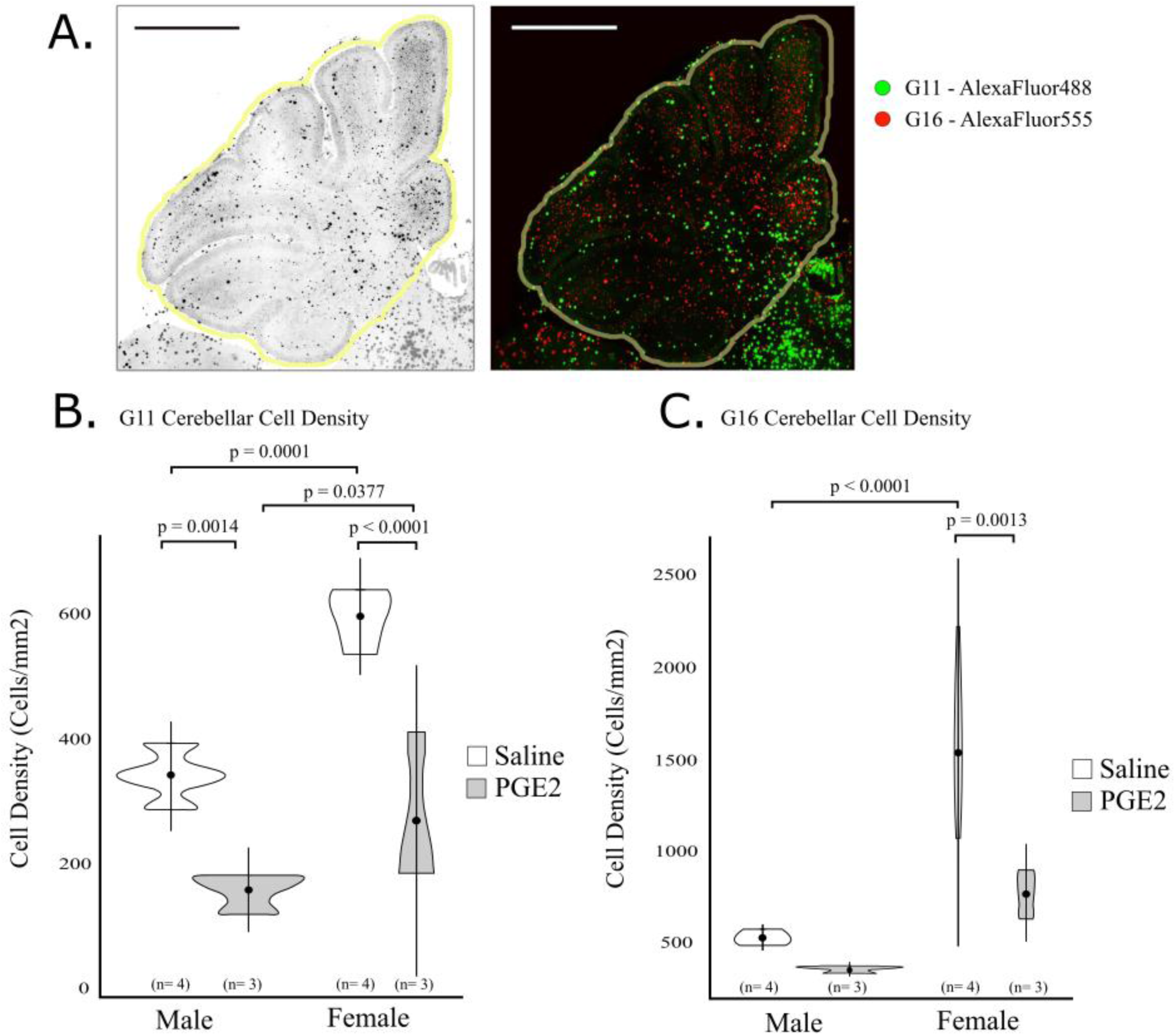
G11 and G16 Cohort-labelled cell densities in the cerebellum of PGE2-exposed mice. (A) CldU and IdU immunolabelled G11 (red) and G16 (green) cells in the cerebellum at PN8. (B) Decreased cell density of G11-labelled cells in males and females. In the Saline controls and PGE2-exposed groups, cerebellar G11 cell density was greater in females than males. (C) Increased cell density of G16-labelled cells observed in females but not males. Cerebellar G16 cell density was greater in Saline females than males. Means represent at least 3 independent animals (shown as n) for each experimental group from 3 separate litters. Data are presented as mean ±SD, with significant differences shown above.

A linear mixed model was also used to fit G16 cell density within the cerebellum. Litter was assigned as a random control, and the fixed effects of condition and sex, as well as the interaction between condition and sex were examined. While the interaction between condition and sex (t(14.0) = - 2.208, p = 0.0444) and the main effect of sex (t(14.0) = 5.693, p < 0.001) were significant, no significance of the condition main effect was found (t(14.0) = -0.892, p = 0.3875). Given the significance of the interaction between condition and sex and sex alone, we performed further pairwise comparisons. No significant difference was found in G16 cell density between PGE2-exposed males and Saline males (Fig 1C, p = 0.3875, Saline = 526.633, PGE2 = 356.428). However, a significantly lower cell density was found within the cerebellums of PGE2-exposed females than in Saline females (Fig 1C, p = 0.0013, Saline = 1532.498, PGE2 = 766.310). While the cell density of Saline males was significantly lower than in Saline females (Fig 1C, p < 0.001, M = 526.633, F = 1532.498), there was no significant difference between PGE2-exposed males and females (Fig 1C, p = 0.0642, M = 356.428, F = 766.310).

To summarize, we observed a significant decrease in cell density within the cerebellum of PGE2-exposed mice depending on the origin of the cells. At G11, PGE2 reduced cell density in both males and females, and at G16 there was a female-specific reduction in cell density within the cerebellum.

### Dendrite length

Average length of primary dendrites was measured from cells within the mature cerebellum at PN30 (Fig 2A and methods) and compared between Saline control and PGE2-exposed male and female mice. Dendrite length was fit using a linear mixed effects model, with litter assigned as a random control. The fixed effects of condition and sex, and the interaction between condition and sex were examined (Fig 2B). We observed no significant effect of the interaction between condition and sex (t(60.0) = -0.708, p = 0.4820), or condition (t(60.0) = 1.474, p = 0.1458), but a significant effect of sex (t(60.0) = 2.160, p = 0.0348) on dendrite length. Given the significant effect of sex, we performed further pairwise comparisons to examine sex-differences within conditions. Cells within the cerebellums of Saline females, were significantly shorter dendrites than in Saline males (Fig 2B, p = 0.0107, M = 23.346, F = 38.017). Interestingly, there was no significant difference between PGE2-exposed male and female mice (Fig 2B, p = 0.2508, M = 33.355, F = 41.230) suggesting a loss of the innate sex difference seen in Saline mice.

**Figure 2.**
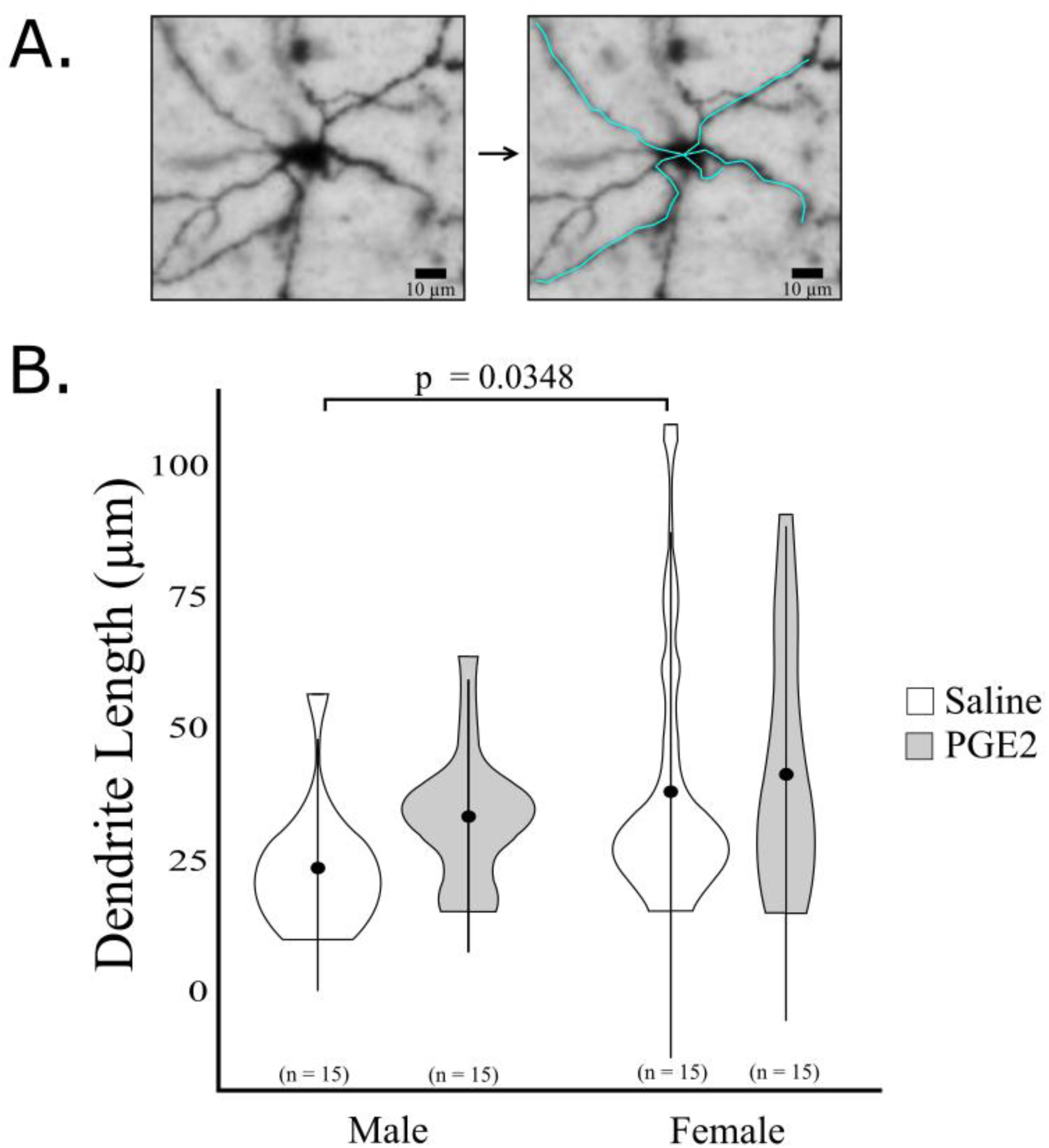
Average dendrite length of cerebellar cells inPN30 Mice. Average length of primary dendrites was measured from cells of Saline controls and PGE2-exposed male and female mice at PN30 from at least 3 separate litters pr condition. Length data are presented as mean ± SD. *n* = 15 cells per condition.

### Dendritic arborization

Dendritic arborization is used to provide information about the ability of a neuron to integrate synaptic potentials [92]. The degree of branching was obtained using sholl analysis to determine the number of intersections made with concentric circles every 10µm from the center of the cell soma (Fig 3A) [93]. We have previously found an increase in dendritic arborization in COX-2^-^-KI mice closer to soma [56]. Here we examined the number of intersections in the PGE2-exposed mice using linear mixed modelling (Fig 3B). The model of *best-fit* was determined to exclude sex as a factor and as such sex was removed from dendritic arborization analysis. Litter was assigned as random control and we examined the fixed effects of condition, and the distance from the soma, as well as the interaction between condition and distance. The significant overall effect of condition (t(715) = -6.289, p < 0.001), prompted us to further investigate pair-wise comparisons at each distance between Saline controls and PGE2-exposed mice.

**Figure 3.**
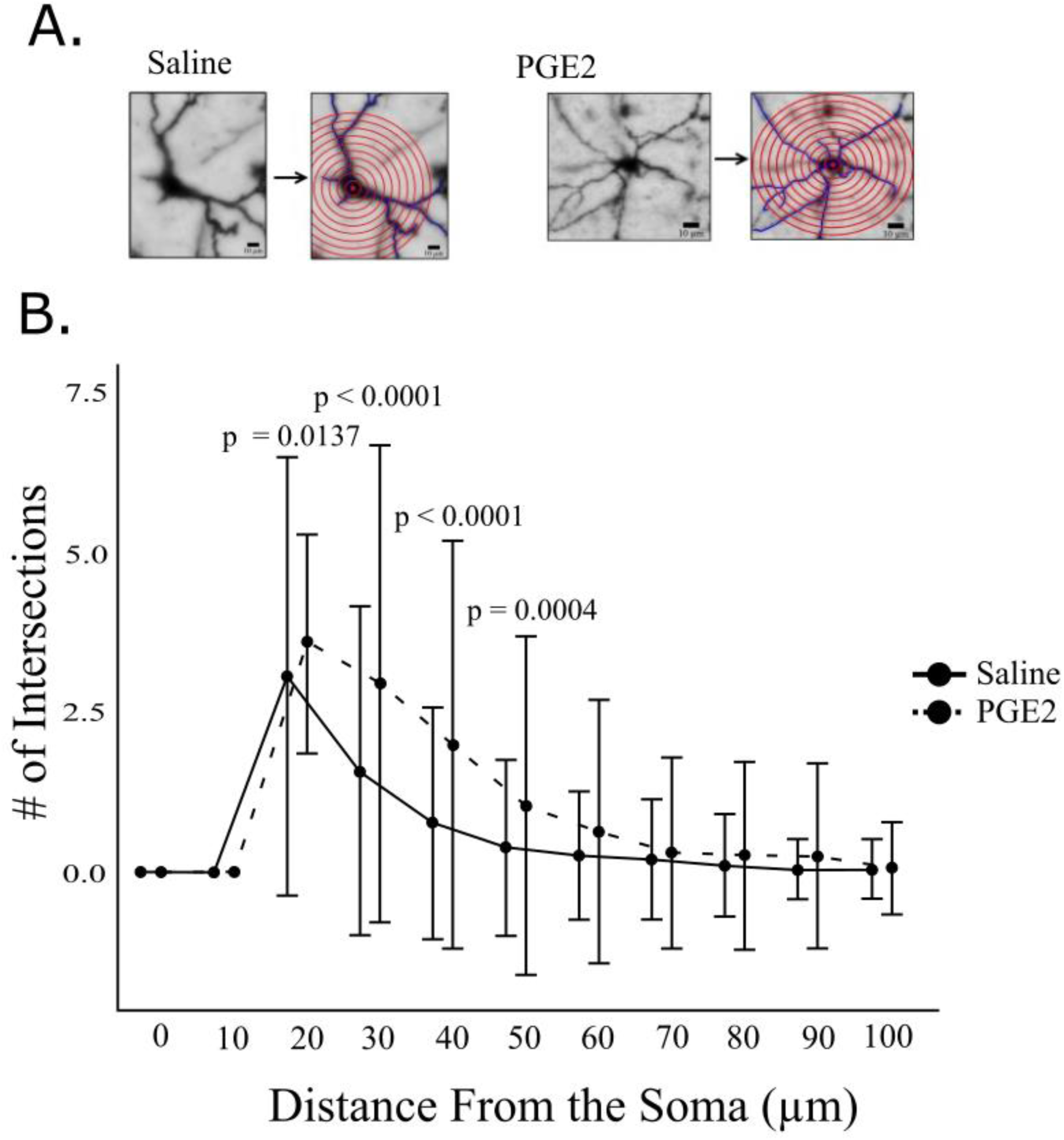
Sholl analysis in the cerebellum of PN30 mice. A.The average number of intersections of dendrites with concentric circles drawn at 10 µm intervals from the cell soma shown here in representative cells from Saline controls and PGE2-exposed. B. Data are presented as mean as mean ± SD (males and female combined). *n* = 30 cells per condition from male and female mice derived from at least 3 separate litters per condition.

Dendritic arborization was compared between Saline controls and PGE2-exposed mice at 10 µm increments from 0 µm to 100 µm (Fig 3B). There was no significant differences in dendritic arborization between Saline controls and PGE2-exposed mice at 0 (t(715) = 0, p = 1.000, Saline = 0.000, PGE2 = 0.000), and 10 µm (t(715) = 0, p = 1.000, Saline = 0.000, PGE2 = 0.000) distances. However, we observed a significantly increased number of intersections in PGE2-exposed mice at 20 µm (t(715) = -2.472, p = 0.0137, Saline = 3.057, PGE2 = 3.600), 30 µm (t(715) = -6.202, p <0.001, Saline = 1.571, PGE2 = 2.933),, 40 µm (t(715) = -5.594, p < 0.001, Saline = 0.771, PGE2 = 2.000), and 50 µm (t(715) =-2.884, p = 0.0040, Saline = 0.400, PGE2 = 1.033) distances. No significant differences between the conditions were observed at 60 µm (t(715) = -1.713, p = 0.0871, Saline = 0.257, PGE2 = 0.633), 70 µm (t(715) = -4.554, p = 0.6490, Saline = 0.200, PGE2 = 0.300), 80 µm (t(715) = -0.6939, p = 0.4870, Saline = 0.114, PGE2 = 0.267), 90 µm (t(715) = -0.802, p = 0.4226, Saline = 0.057, PGE2 = 0.233), and 100 µm (t(715) = -0.043, p = 0.9654, Saline = 0.057, PGE2 = 0.067) distances from the center of the cell soma.

In summary, there was greater dendritic arborization within the cells of the cerebellums in PGE2-exposed mice at distances closer to the some particularly between the 20-50µm range.

### Dendritic looping

We have previously demonstrated *in vitro* that exposure to PGE2 during the differentiation of NE-4C stem cell significantly increased the proportion of dendrites that formed loops [86]. We have also found an increase in the odds of observing dendritic loops in the cerebellum of COX-2^-^-KI mice [39, 56]. Here we investigate the odds of observing dendritic loops in the cerebellum of PGE2-exposed mice. The greatest exterior angle of all dendrites found was measured and dendrites with an exterior angle exceeding 270° were classified as loop forming. (Figure 4, methods). With this distinction made, there were differences in the distribution of dendrites forming loops with higher proportion of looping dendrites exceeding 270° in PGE2-exposed mice compared to Saline control mice. To further investigate these differences, and given the categorical nature of the loop classification, we fit loop classifications using a binomial logistic regression. We examined the odds that any given dendrite would form a dendritic loop given its condition (Saline controls or PGE2-exposed) and sex (Table 1).

**Figure 4.**
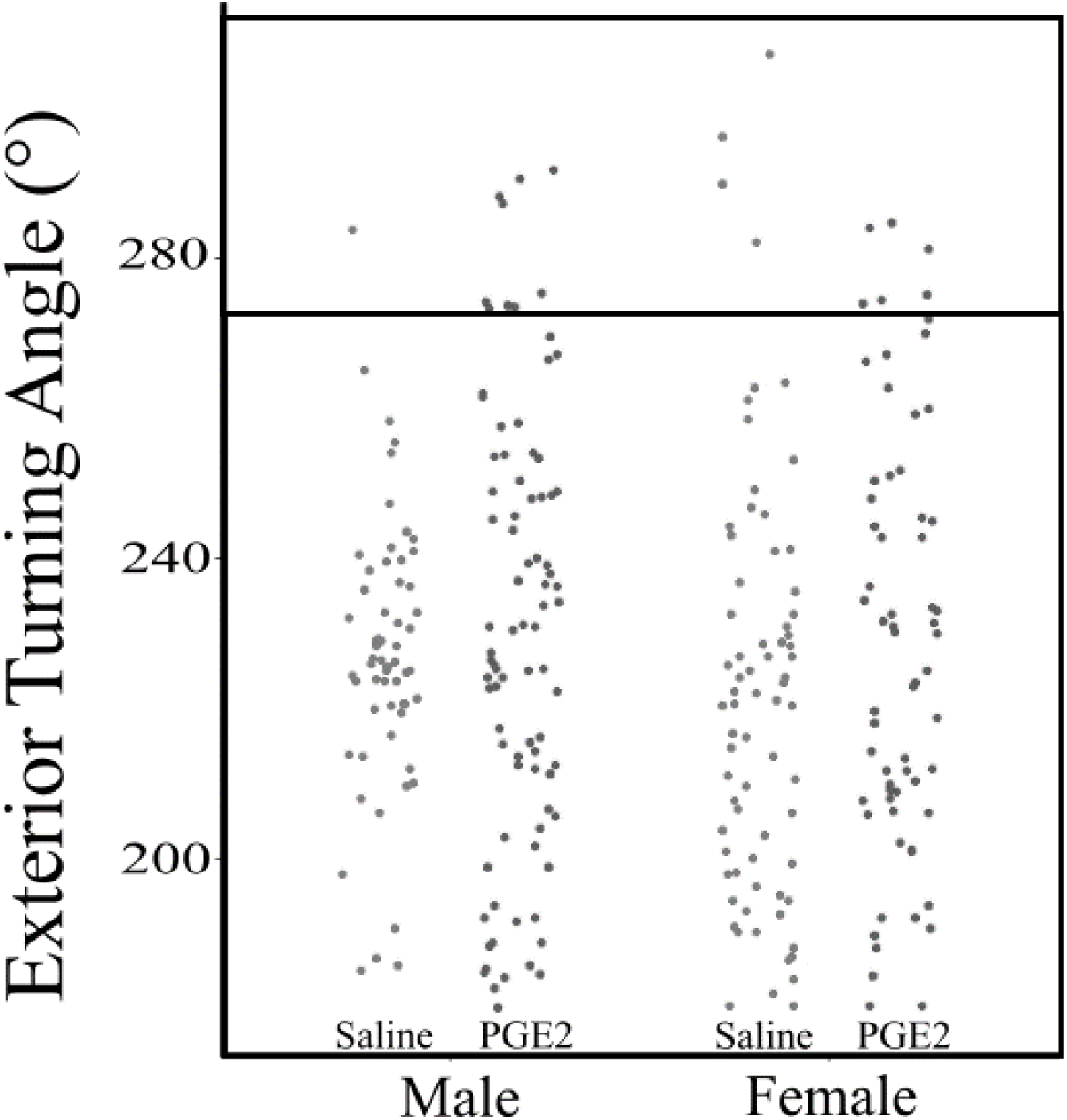
Distribution of turning angles of dendrites in the cerebellum. The greatest exterior angle was measured for dendrites within the PN30 cerebellum of Saline control and PGE2-exposed male and female mice from at least 3 separate litters per condition. Dendrites with a turning angle greater than 270° were classified as axonal loops. Each dot represents individual dendrite.

**Table 1.**
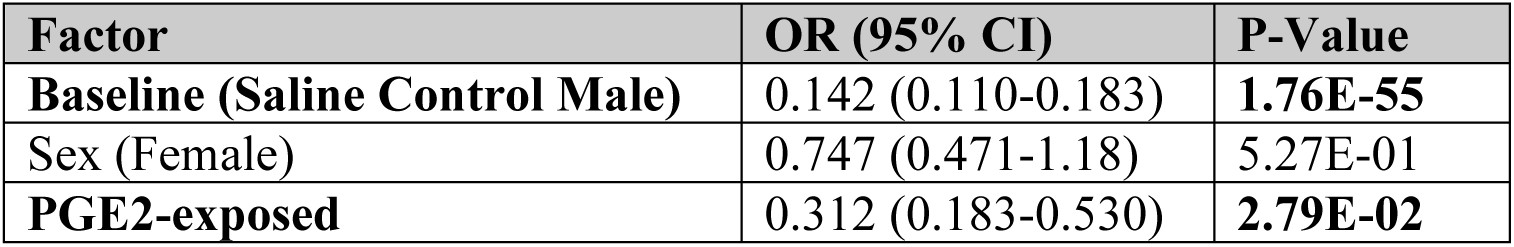
Odds of observing dendritic loops in the cerebellum of PGE2-exposed offspring. Likelihood of a given dendrite being classified as looping was determined for Golgi-COX stained cerebellar neurons of mice at PN30. Odds ratios (95% confidence intervals) are given for the baseline (Saline control male) as well as the factors of sex and PGE2-exposure. At least 10 cells from 3 animals per condition from 3 separate litters were used (n = 30 per condition).

We used multinomial logistic regression to determine the odds that our baseline intercept (Saline control male) would form dendritic loops (Methods). We found a significant odds ratio (OR) of 0.142 in the saline control male intercept (p < 0.001), signifying that it was less likely to observe a looping dendrite than a non-looping dendrite in these saline control males (Table 1). Examining the sex factor (if the animal was female), we determined that sex did not affect the likelihood of observing dendritic loops from what was observed in Saline control males (OR = 0.747, p = 0.5270). However, the condition factor (if the animal was exposed to PGE2) significantly increased the odds of observing a dendritic loop (OR = 0.312, p = 0.0279).

Overall, while sex did not have a significant effect, we found that PGE2-exposed offspring have increased odds of observing dendrites that are considered looping.

### Dendritic spine density

Studies in other labs have shown that local injection of PGE2 within the cerebellum during critical time points in its postnatal development affects dendritic spine density. For example, injection of PGE2 at PN10 and PN12 into the cerebellum of rats inhibited dendritic spine formation [94]. In this study we examined the effect of prenatal exposure to PGE2 at G11 on dendritic spine density and morphology within the mature cerebellum at PN30 mice offspring (Fig 5A).

**Figure 5.**
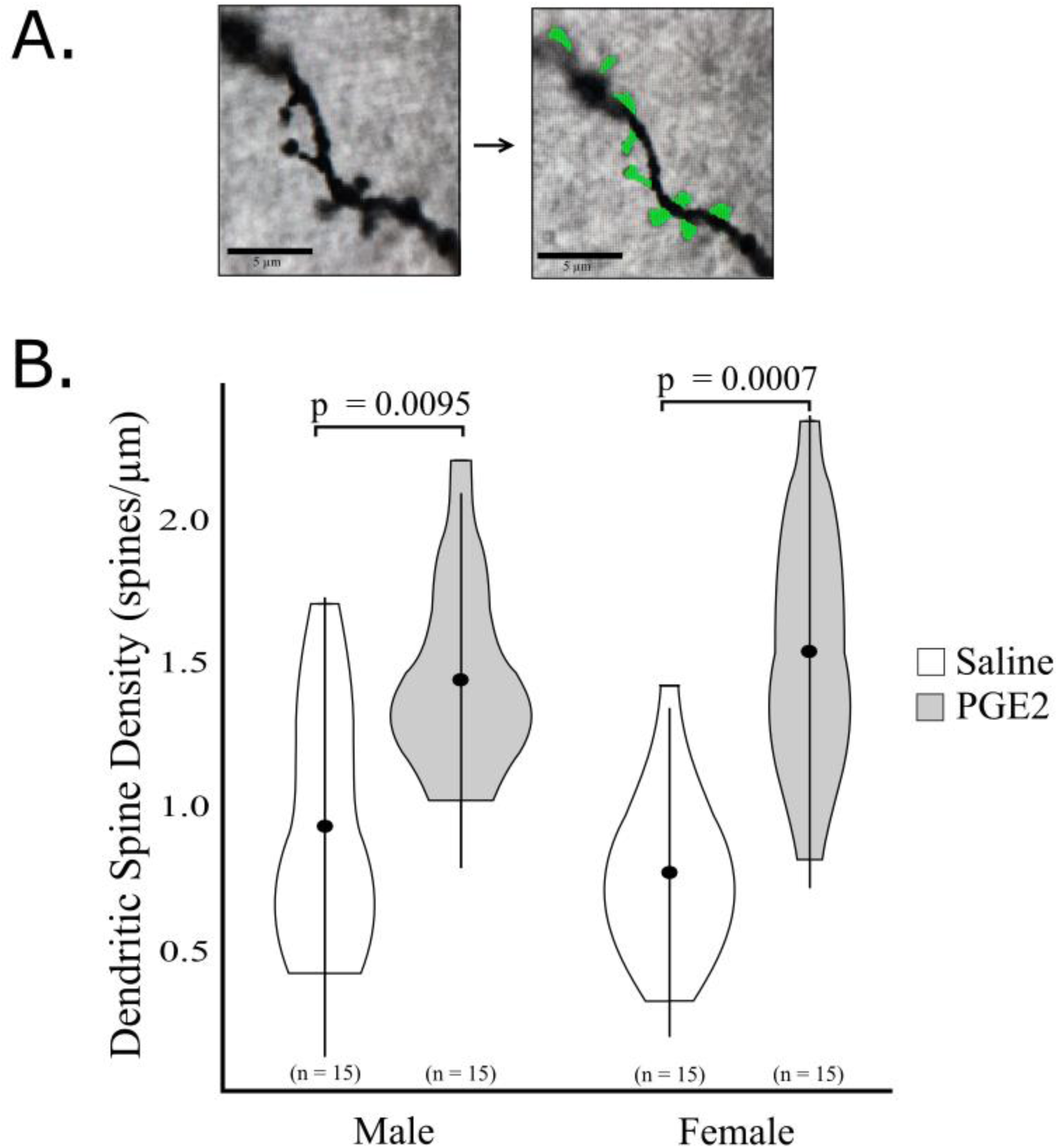
Dendritic spine density in the cerebellum of P30 Mice. Dendritic spines were detected by AI Software (Nikon). (A) A representative Golgi-Cox staining of dendrites in cells of the cerebellum with detected spine shapes shown in green (*right* image). (B) The number of spines per µm of dendrite length were measured from cerebellums of Saline control and PGE2-exposed male and female offspring from at least 3 different litters per condition. Data are presented as mean ± SD. *n* = 15 number of dendrites per condition.

First, dendritic spine density (spines/µm) was fit using a linear mixed effects model, with litter assigned as a random control. The fixed effects of condition and sex, and the interaction between condition and sex were examined (Fig 5B). While we saw no significant effect of the interaction between condition and sex (t(54.0) = 1.572, p = 0.1218), or the main effect of sex (t(54.0) = -1.360, p = 0.1794), there was a significant effect of condition (t(10.7) = 3.151, p = 0.0095) on dendritic spine density. Given the significant effect of condition, we examined pairwise comparisons between PGE2-exposed mice and their sex-matched Saline controls. Exposure to PGE2 significantly increased dendritic spine density in both males (p = 0.0095, Saline = 0.9249, PGE2 = 1.4368), and females (p = 0.0007, Saline = 0.7666, PGE2 = 1.537). In summary, PGE2-exposed mice showed an increase in dendritic spine density compared to Saline control mice regardless of gender.

### Dendritic spine morphology

Dendritic spines change shape as they mature, progressing from thin, to stubby to mushroom [95–97]. These changes in dendritic spine morphology correspond to their function of the brain and influence plasticity of the neuron, and ultimately the brain as a whole [98, 99]. We have previously shown that COX-2^-^-KI male mice were less likely and COX-2^-^-KI females were more likely to have mature spines compare to the Saline control counterparts [39, 56]. Here, we investigate the effect of prenatal PGE2 exposure on dendritic spine morphology in male and female offspring at PN30. We investigated the relative odds that thin or stubby shaped spines would be observed compared to mushroom shaped spines in each condition. A multinomial logistic regression was used to fit the data and the likelihood that any given spine was mushroom shaped or was thin or stubby shaped was determined by condition. Litter was assigned as a random effect to control for potential litter-bias. An odds-ratio (OR) above 1 for a shape within a condition signified that there was a higher likelihood of observing that shape compared to a mushroom shaped spine.

We assigned Saline male controls as our baseline intercept for comparisons. First, we established that in Saline male controls, there was a higher likelihood of observing mushroom shaped spines compared to both stubby (Table 2, OR = 0.782, p <0.001), and thin (OR = 0.121, p < 0.001) shaped spines. Next, we found that compared to Saline control males, Saline female controls were more likely to see stubby shaped spines than mushroom shaped spines (OR = 1.427, p < 0.001) but there was no difference for the likelihood of observing thin shaped spines (OR = 1.400, p < 0.001).

**Table 2:** Odds of observing dendritic spine shapes in the cerebellum of PGE2-exposed offspring. The likelihood of a given dendritic spine being mushroom shaped, or thin or stubby shaped was determined for dendritic spines of Golgi-COX-stained cerebellar neurons of male and female PGE2-exposed and Saline control mice at PN30. Odds ratios (95% confidence intervals) are given for the likelihood of each shape in each condition. All dendritic spines were measured from at least 10 cells from 3 animals per condition from 3 separate litters.

| Factor | Shape | OR (95% CI) | P-Value | Odds of Mature spines (Compared to Saline Control Male) |
| --- | --- | --- | --- | --- |
| Saline Control Male | **Stubby** | 0.782 (0.730– 0.838) | **<0.001** | - |
|  | **Thin** | 0.121 (0.105– 0.139) | **<0.001** |  |
| Saline Control Female | **Stubby** | 1.427(1.295 – 1.574) | **<0.001** | Decreased |
|  | Thin | 1.400 (1.156 – 1.697) | 0.0795 |  |
| PGE2-exposed Male | **Stubby** | 0.810 (0.738 – 0.889) | **0.0145** | Increased |
|  | **Thin** | 0.596 (0.486 – 0.732) | **0.0116** |  |
| PGE2-exposed Female | **Stubby** | 0.646 (0.568 – 0.735) | **< 0.001** | Increased |
|  | Thin | 1.346 (1.023 – 1.773) | 0.2794 |  |

In PGE2-exposed males we found an increased likelihood of observing mushroom-shaped spines compared to stubby (OR = 0.810, p = 0.0145), or thin (OR = 0.596, p = 0.0116) shaped spines relative to the likelihood in Saline male mice. Similarly, in PGE2-exposed females had an increased likelihood of observing mushroom-shaped spines compared to stubby shaped spines (OR = 0.646, p <0.001) but there was no difference in the likelihood of observing thin shaped spines (OR = 1.346, p = 0.2794) compared to the likelihood in Saline male controls.

In summary, we observed an innate sex difference in dendritic spine morphology between Saline male and female control mice, with females being more likely to observe immature stubby shaped spines and males with more mature mushroom spines. However, PGE2-exposure increases the odds of observing mature mushroom shaped spines in males and females.

### Adhesive sticker test

We have previously established that that the single maternal exposure to PGE2 results in autism-related behaviours in offspring postnatally including deficits in social interaction, anxious behaviours, and repetitive or restrictive behaviours [38, 39, 56]. The goal of this study is to evaluate whether PGE2-exposed mice display specific cerebellum-related behaviours as a consequence of the morphological changes described. First, we use the *adhesive sticker test* to measure sensory motor coordination in mice as they attempted to remove a small adhesive sticker from just above their noses [78–80] (Fig 6A). To verify that there was not a confounding effect of anxiety on differences in motor coordination, we measured the time it took the mouse to make first contact with the adhesive sticker. A linear mixed effect model was used to fit the time until mice made first contact with the adhesive sticker (Fig 6B). Litter was assigned as a random control and the fixed effects of condition and sex, as well as the interaction between condition and sex were examined. We observed no significance in sex, condition, or the interaction between sex and condition indicating that any differences in the adhesive sticker test would not be due to anxious behaviour.

**Figure 6.**
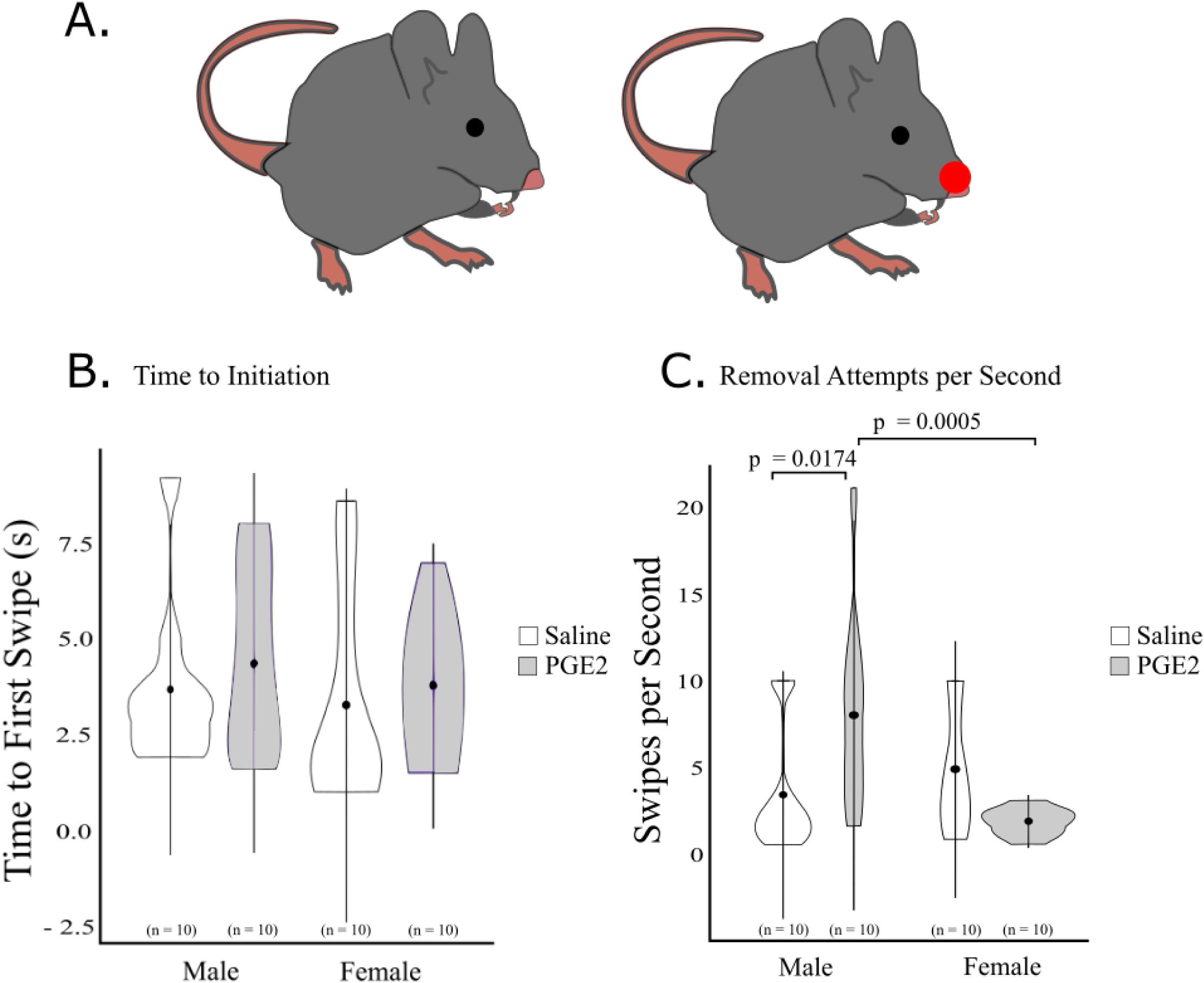
Adhesive Sticker Test of motor coordination. A. A small adhesive sticker was placed just above the nose of Saline control and PGE2-exposed mice on PN30. The time until the mouse made first contact with the adhesive sticker and the number of attempts to remove the sticker were per second were recorded. B. No significant differences were observed between conditions in the time until the first removal attempt was made. C. PGE2 exposed males made significantly more attempts per second than both C57 controls and PGE2 exposed females. *n* = 10 mice per condition, derived from at least 4 separate litters per condition Data are presented as mean ±SD, with significant differences shown above.

Next, we measured performance in the *adhesive sticker test* as the number of swipes each mouse made, which was corrected with the time taken by the mouse to remove the sticker (see methods). Swipes per second was fit using a linear mixed effects model, with litter assigned as a random control. The fixed effects of condition and sex, and the interaction between condition and sex were examined (Fig 5C). We observed a significant interaction between condition and sex (t(38.4) = - 3.297, p = 0.0021) and the main effect of sex (t(37.6) = 3.764, p = 0.0006), but no significant effect of condition alone (t(11.2) = 1.79, p = 0.1010). Given the significance of the interaction and the main effect of sex, we performed further pairwise comparisons. There was a statistically significant increase in the number of swipes per second in PGE2-exposed males compared to Saline male controls (Fig 6C, p = 0.0174, Saline = 3.421, PGE2 = 8.029). However, there was no significant difference in swipes per second between PGE2-exposed and Saline control female mice (Fig 6C, p =0.1010, Saline = 4.876, PGE2 = 1.913). While there was no significant difference in swipes per second in Saline male and female control mice (Fig 6C, p = 0.3822, M = 3.421, F = 4.876), PGE2-exposed males had a significantly higher number of swipes per second than PGE2-matched females (Fig 6C, p < 0.001, M = 8.029, F = 1.913).

In summary, we observed an PGE2-male specific increase in swipes per second, demonstrating reduced motor coordination in these mice.

### Grid walking test

*The grid walking test* further tests sensory motor coordination of mice as they attempted to cross a mesh grid. It measures the number of limb slips each mouse made through the grid and the time taken by each mouse to cross the grid [81, 82] (Fig 7A). We first examined the number of limb slips made by each mouse as they crossed the grid (Fig 7B). Number of slips through the grid was fit using a linear mixed effects model, with litter assigned as a random control. The fixed effects of condition and sex, and the interaction between condition and sex were examined (Fig 7B). While the interaction between condition and sex was significant (t(95.5) = -3.016, p = 0.0033), we saw no significant effect of the condition (t(11.2) = 0.872, p = 0.4017), or sex (t(90.6) = -0.065, p = 0.9484) main effects on number of slips. Given the significance of the interaction between condition and sex, further pairwise comparisons were performed. We saw no significant difference between PGE2-exposed mice and sex matched Saline controls in males (Fig 7B, p = 0.4017, Saline = 4.1, PGE2 = 4.718), or females (Fig 7B, p = 0.3236, Saline = 4.289, PGE2 = 3.317). While there was no difference in the number of slips through the grid between Saline male and female controls (Fig 7B, p = 0.3878, M = 4.1, F = 4.289), PGE2-exposed males made significantly more slips through the grid than PGE2-exposed females (Fig 7B, p = 0.0001, M = 4.718, F = 4.289).

**Figure 7.**
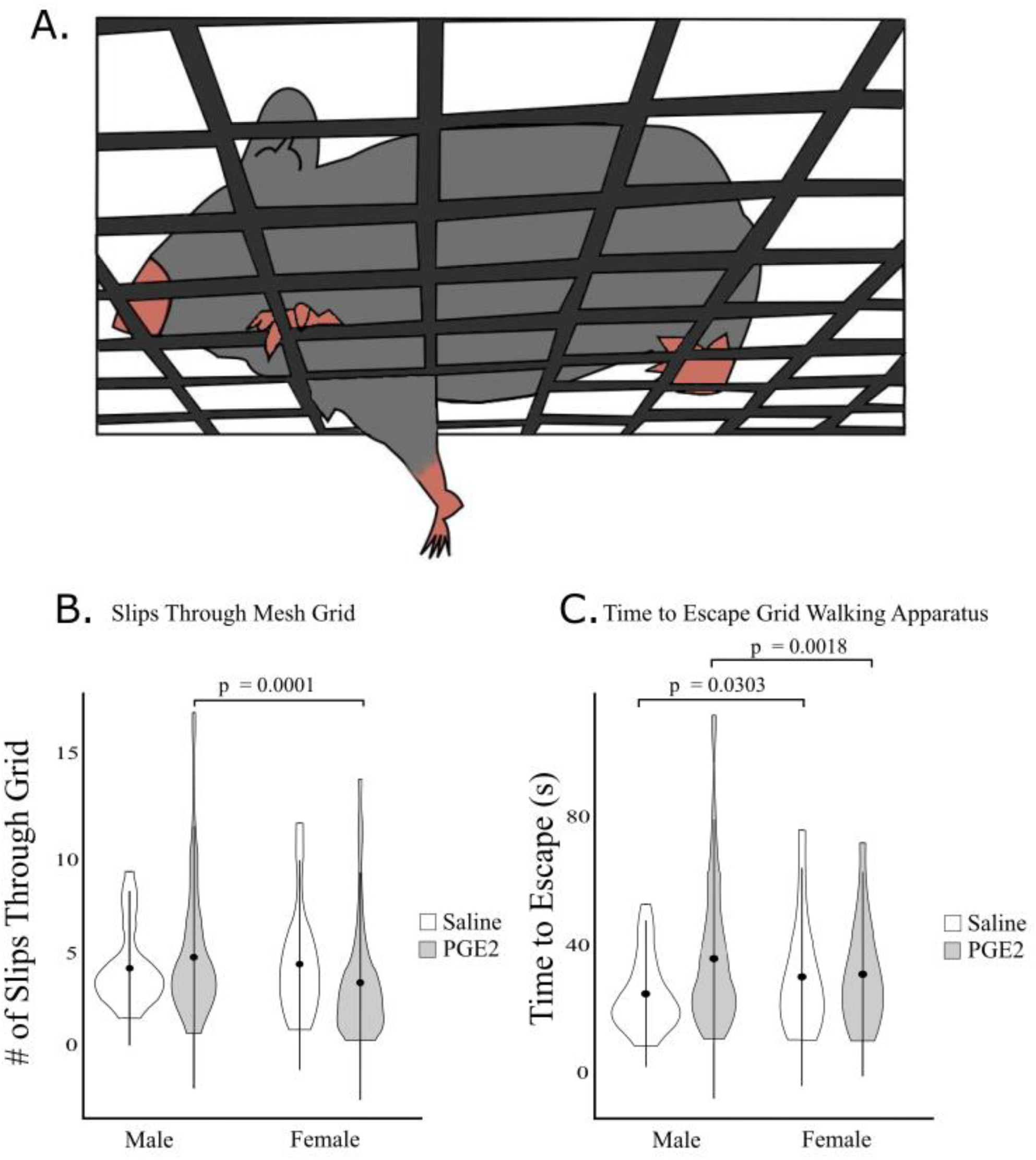
Grid Walking Test of motor coordination. A. Mice were allowed to cross an enclosed mesh grid on PN30. The number of limb slips of each mouse moving through the grid as well as the time it took each mouse to escape the grid were recorded. B. PGE2 males slipped significantly more than PGE2 females. C. Saline male controls were quicker to escape the grid than Saline female controls, but PGE2-exposed males were slower to escape than PGE2-exposed females. Means represent 10 animals for each experimental group taken from at least 4 separate litters per condition. Data are presented as mean ±SD, with significant differences shown above.

The time taken by each mouse to escape the grid was also examined (Fig 7C). Time to escape was fit using a linear mixed effects model, with litter assigned as a random control. The fixed effects of condition and sex, and the interaction between condition and sex were examined. We observed a significant effect of the interaction between condition and sex (t(95.0) = -3.872, p = 0.0002), and the main effect of sex (t(90.8) = 2.200, p = 0.0303), but no significant effect of the condition main effect (t(11.1) = 1.679, p = 0.1211). Given the significant interaction between condition and effect and the significant effect of sex, we performed further pairwise comparisons. We saw no significant differences between PGE2-exposed and Saline control males (Fig 7C, p = 0.1211, Saline = 24.642, PGE2 = 35.605), or females (Fig 7C, p = 0.5487, Saline = 29.963, PGE2 = 31.179). We observed opposite effects between sexes within Saline controls and PGE2-exposed mice. Though Saline male controls escaped significantly faster than Saline female controls (Fig 7C, p = 0.0303, M = 24.642, F = 29.963), PGE2-exposed males were slower to escape than PGE2-exposed females (Fig 7C, p = 0.0018, M = 35.605, F = 31.179).

In summary we observed a sex-dependent effect of PGE2 on motor coordination in the grid walking test. In number of slips a sex difference was present in the PGE2-exposed mice that was not seen in Saline control mice. In time to escape there were sex differences within Saline controls with female being slower and within PGE2-exposed mice with male being slower.

### Cylinder test

The cylinder test measures exploratory motor behaviour as mice explore a tall glass cylinder [83–85]. In this study, we use the cylinder test to measure the number of forelimb touches to the side of the cylinder, the total number of rears, and the total number of steps (Fig 8A). Forelimb touches were quantified as the number of times the mouse would touch the side of the cylinder with either of their forelimbs. Forelimb touches were fit using a linear mixed effects model, with litter assigned as a random control. The fixed effects of condition and sex, and the interaction between condition and sex were examined (Fig 8B). While we observed no significant effects of the interaction between condition and sex (t(110.1) = -1.138, p = 0.2578), or the condition main effect (t(0.6) = 1.708, p = 0.5414), there was a significant effect of sex (t(97.7) = 2.662, p = 0.0091). Given the significant effect of the sex condition, we examined pairwise comparisons between sex within each condition. There were significantly fewer forelimb touches by Saline male control mice than Saline female control mice (Fig 8B, p = 0.0091, M = 5.236, F = 9.008). However, there were no sex-differences in forelimb touches between PGE2-exposed males and females (Fig 8B, p = 0.285, M = 5.592, F = 7.110).

**Figure 8.**
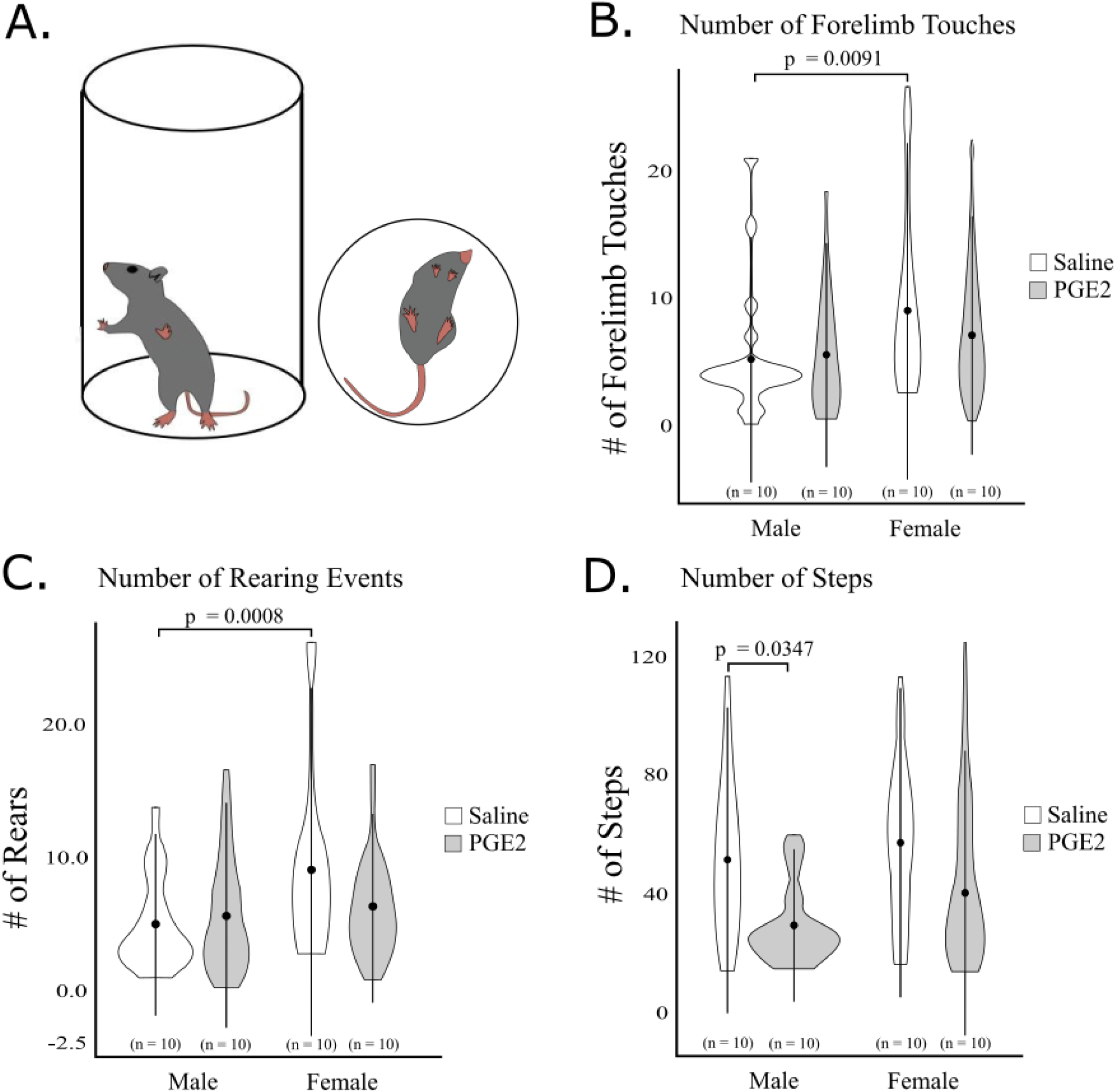
Cylinder Test of motor coordination. A. Saline control and PGE2-exposed mice on PN30 were placed into a long clear cylinder (*side* and *top* view). The number of forelimb touches to the cylinder, the number of rears, and the number of steps were recorded. B. Saline female controls made significantly more touches than Saline male controls with no difference between PGE2-exposed mice. C. Saline female controls reared more than Saline male controls again with no difference between PGE2-exposed mice. D. PGE2-exposed males made significantly less steps than Saline male controls. Means represent 10 animals for each experimental group from at least 4 separate litters. Data are presented as mean ±SD, with significant differences shown above.

Rears were measured whenever the mouse would rise up of its forelimbs onto its hind limbs. Rears were fit using a linear mixed effects model, with litter assigned as a random control. The fixed effects of condition and sex, and the interaction between condition and sex were examined (Fig 8C). We observed a significant effect of the interaction between condition and sex (t(104.0) = -2.082, p = 0.0398), and sex (t(97.4) = 3.476, p = 0.0008), but no significant effect of condition was observed (t(14.9) = 0.900, p = 0.3825). Given the significant effects of the interaction and the sex main effect, we performed further pairwise comparisons. We observed no significant difference in the number of rears between Saline and PGE2-exposed males (Fig 8C, p = 0.3825, Saline = 4.952, PGE2 = 5.675), or between Saline and PGE2-exposed females (p = 0.2798, Saline = 8.942, PGE2 = 6.196). There were sex-differences between conditions. Saline male controls reared significantly less than Saline female controls (Fig 8C, p = 0.0008, M = 4.952, F = 8.942). However, this sex-difference was no longer observed between PGE2-exposed males and females (Fig 8C, p = 0.6008, M = 5.675, F = 6.196).

We also measured the number of steps taken by each mouse while in the cylinder. Steps were fit using a linear mixed effects model, with litter assigned as a random control. The fixed effects of condition and sex, and the interaction between condition and sex were examined (Fig 8D). While there was no significant interaction between condition and sex (t(100.4) = 1.559, p = 0.1221), or a significant effect of sex (t(93.0) = 0.076, p = 0.9396), we observed a significant effect of condition (t(13.6) = - 2.346, p = 0.0347) on number of steps. Given the significant effect of condition, we performed further pairwise analysis on differences based on condition. We saw a significant reduction in the number of steps taken by PGE2-exposed males compared to Saline male controls (Fig 8D, p = 0.0347, Saline = 51.138, PGE2 = 29.233). However, there was no significant difference in number of steps between PGE2-exposed and Saline female controls (Fig 8D, p = 0.3607, Saline = 57.013, PGE2 = 40.082).

In summary, we found changes in motor function in PGE2-exposed mice including loss of the innate sex difference observed in Saline control mice for forelimb touches and number of rears, and a reduction in the number of steps taken, which was specific to males.

### Cerebellar protein expression

In this study, we have found that prenatal exposure to PGE2 affects density of migrating cells and the morphology of dendrites and dendritic spines within the cerebellum of male and female offspring. Dendritic morphology is normally regulated by cytoskeletal dynamics [100–102]. Our previous studies in differentiating NE-4C stem cells exposed to PGE2 [66] and COX-2^-^-KI mice [74] showed abnormal expression of β-actin and the actin-binding protein spinophilin highly abundant in dendritic spines.

Here, we examine whether the expression levels of spinophilin, and β-actin are also affected in the cerebellum of PGE2-exposed offspring at P30. In addition, we analyze the level of N-Cadherin, a protein that promotes dendritic outgrowth and stabilization of mature synapses [103–107]. For each protein linear mixed modelling was used to fit the fold change relative to the expression of in Saline male controls for each protein across 3 separate western blot runs (Fig 9). To avoid confounding by differences between technical replicates, we assigned technical replicates as a random effect.

**Figure 9.**
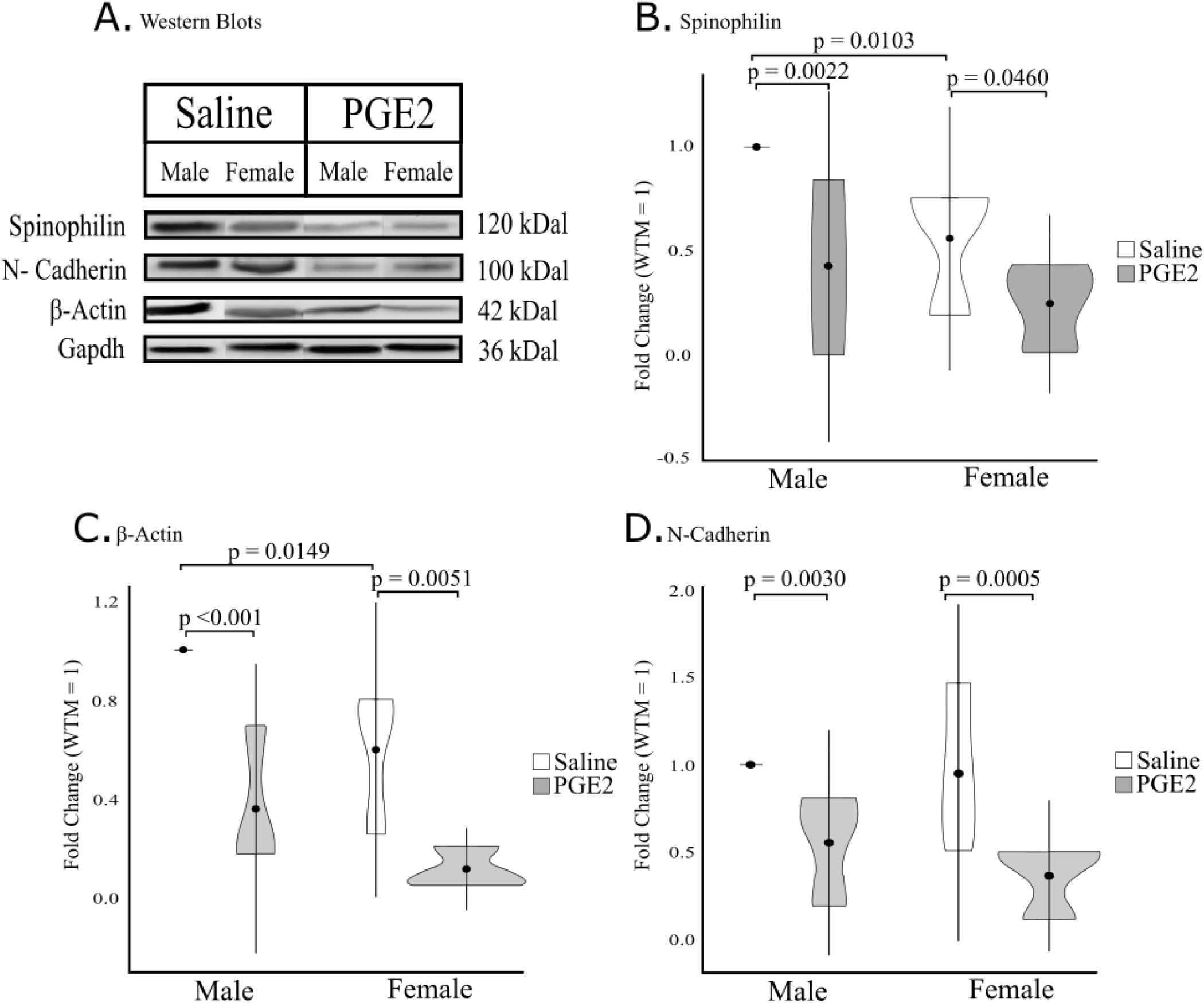
Protein expression of Spinophilin, β-Actin and N-Cadherin within P30 mice. A. Cerebellar isolates of Saline control and PGE2-exposed male and female mice were quantified for the expression levels of Spinophilin (B), β-Actin (B) and N-Cadherin (D). Protein expression was normalized to the house keeping gene GAPDH and then quantified relative to the expression in Saline male controls. All samples are pooled from 10 animals derived from at least 4 separate litters. Data are presented as mean fold change ± SD. Each graph represents is an average of 3 individual Western blot runs per condition.

Condition, sex and the interaction between condition and sex were tested for spinophilin expression level. While there was no effect of the interaction between condition and sex (t(9.0) = 1.355, p = 0.2084), we found significant effects of both condition (t(9.0) = -4.230, p = 0.0022), and sex (t(9.0) = -3232, p = 0.0103) alone. Given the significant effects of condition, and sex we performed further pairwise comparisons to examine the effects of each. Compared to sex-matched Saline controls, there was a significant reduction in the expression of spinophilin in PGE2-exposed males (Fig 9A and B, p= 0.0022, Saline = 1.000, PGE2 = 0.4241) and females (p = 0.0460, Saline = 0.5600, PGE2 = 0.2449). Within conditions we observed that while the expression of spinophilin was higher in Saline male controls than in female controls (p = 0.0103, M = 1.000, F= 0.5600), there was no significant difference between PGE2-exposed males and females (p = 0.2209, M = 0.4241, F= 0.2449).

Again condition, sex and the interaction between condition and sex were tested for β-actin expression levels. We found no significant effect of the interaction between condition and sex (t(12.0) = 0.807, p = 0.4353), but found significant effects of both condition (t(12.0) = 0.0007, p =), and sex (t(12.0) =, p = 0.0149). Given that the effects of both condition and sex were significant, pairwise comparisons involving condition and sex were further investigated. Similar to spinophilin, the expression of β-actin was significantly lower in PGE2-exposed males (Fig 9A and B, p =0.0007, Saline = 1.000, PGE2 = 0.3556) and females (Fig 9B, p = 0.0051, Saline = 0.5985, PGE2 = 0.1155), compared to sex-matched Saline controls. Moreover, the expression of β-actin was higher in Saline male than female controls (p = 0.0149, M = 1.000, F = 0.5985), but no significant difference was found between PGE2-exposed males and females (Fig 9B, p = 0.1150, M = 0.3556, F = 0.1155).

Lastly, we also tested condition, sex and the interaction between condition and sex for N-cadherin expression levels, with technical replicate assigned as a random effect. While we found no significant effect of the interaction between condition and sex (t(9.0) = -0.556, p = 0.5915), or sex (t(9.0) = -0.277, p = 0.7881), the condition effect was found to be significant (t(9.0) = -2.512, p = 0.0332). As the effect of condition was significant, we examined further pairwise comparisons between conditions within each sex. There was a reduction in N-Cadherin expression in both PGE2-exposed male (Fig 9C, p = 0.0332, Saline = 1.000, PGE2 = 0.5519), and female (Fig 9C, p = 0.0092, Saline = 0.9506, PGE2 = 0.3622) mice compared to sex-matched Saline controls.

In summary, PGE2-exposure resulted in reduced expression levels of β-actin, spinophilin and N-cadherin within the male and female cerebellum, proteins involved in dendritic outgrowth and migration of neuronal cells.

### Behaviour matched protein expression

Our final aim was to assess a potential correlation between the observed behaviour in PGE2-exposed offspring and the expression level of the proteins analysed. Within our behavioural experiments, PGE2 males consistently performed the worst on motor tasks. Thus, from the PGE2-exposed males tested (Male pooled in Fig 9) we selected 3 males from independent litters that performed the best and worst overall across the behavioural tests. Male 1 had the best overall performance (similar to Saline male controls) among all the PGE2 males across tests, Male 2 had the worst scores on the adhesive sticker and cylinder test, and Male 3 had the worst scores on the grid walking test. We compared the cerebellar expression levels of spinophilin, β-actin, and N-Cadherin within these individual males to pooled Saline control and pooled PGE2-exposed male samples (Fig 10A). As above, we used linear mixed modelling to fit the fold-change of each protein expression level relative to Saline male controls across three separate western blot runs (Fig 10B-D).

**Figure 10.**
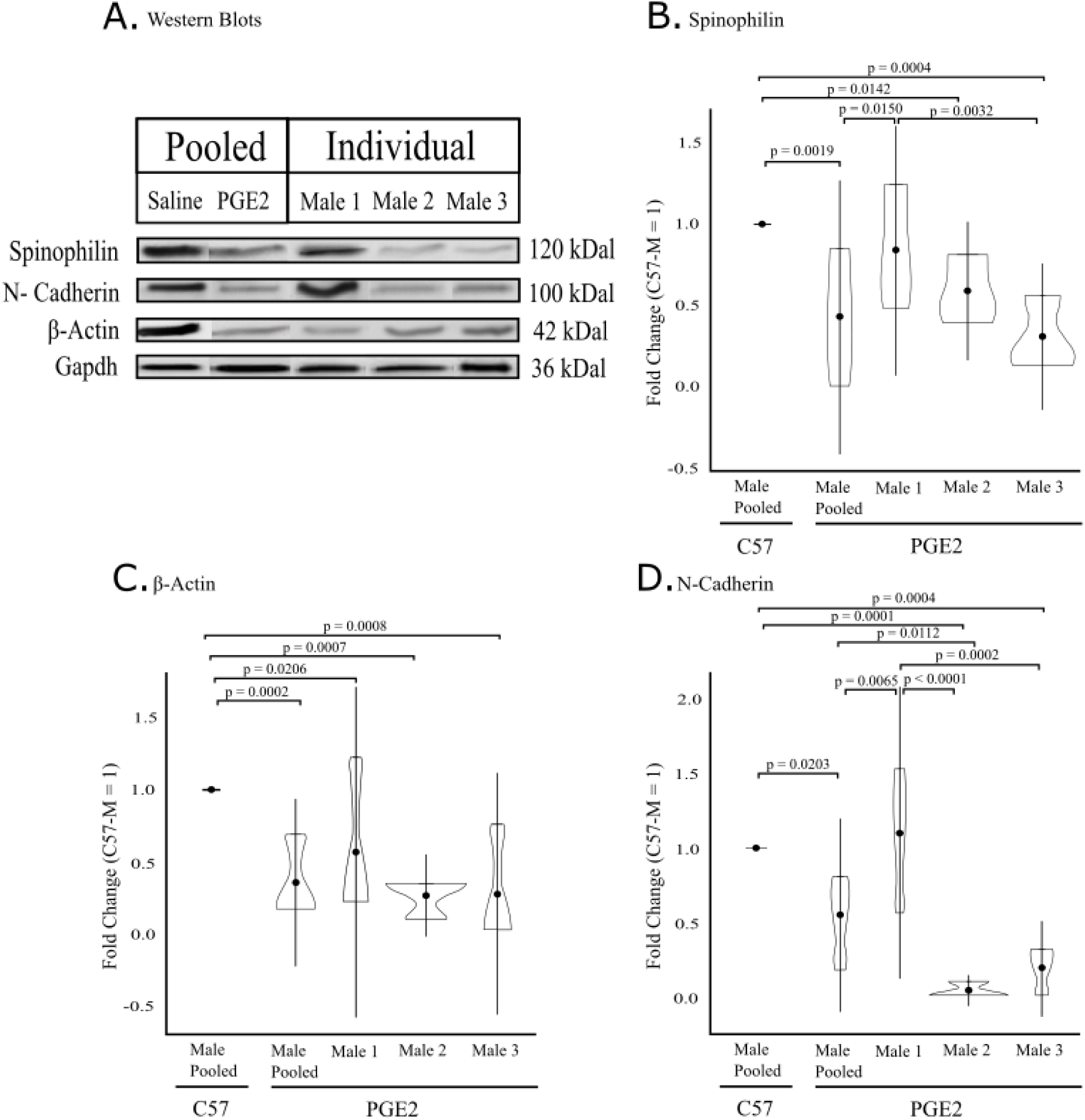
Protein Expression of Spinophilin, β-Actin and N-Cadherin in Saline and PGE2 pooled and individual male mice. Cerebellar samples of pooled Saline controls (C57) and PGE2-exposed male P30 mice and selected PGE2-exposed male mice were analysed for the expression of Spinophilin, β-Actin and N-Cadherin. Protein expression was normalized to the house keeping gene GAPDH and then quantified relative to the expression in the pooled Saline Male controls. Pooled samples derive from 10 animals from at least4 separate litters (as in Figure 9). Male 1 had the best performance in cerebellar behaviour across all PGE2 males, while Males 2 and 3 had the worst performance across all PGE2 males. Data are presented as mean fold change ± SD. Graphs for each protein are representative of 3 individual runs per condition.

First, we compared samples (Saline-Males Pooled, PGE2-males Pooled, Male 1, Male 2, Male 3) to each other using technical replicate as a random effect (Fig 10B). As reported in Figure 9, we found a significant reduction in spinophilin expression in PGE2-males compared to Saline male controls (p = 0.0019, Saline = 1.000, PGE2 = 0.4241). Interestingly, the expression of spinophilin in Male 1 was not significantly different than in Saline male controls (p = 0.2811, Saline = 1.000, Male 1 = 0.8361), but was significantly greater than in PGE2-males (p = 0.0150, PGE2 = 0.4241, Male 1 = 0.8361). In contrast, the expression of spinophilin for Male 2, was significantly lower than Saline male controls (p = 0.0142, Saline = 1.000, Male 2 = 0.5839), but not significantly different than in PGE2-males (p = 0.2927, PGE2 = 0.4241, Male 2 = 0.5839). Similarly, spinophilin expression was lower in Male 3 than Saline male controls (p = 0.0043, Saline = 1.000, Male 3 = 0.3032), but not different than in PGE2-males (p = 0.4217, PGE2 = 0.4241, Male 3 = 0.3032). While we found no differences in spinophilin expression between Male 1 and Male 2 (p = 0.1081, Male 1 = 0.8361, Male 2 = 0.5839), or Male 2 and Male 3 (p = 0.0773, Male 2 = 0.5839, Male 3 = 0.3032), spinophilin expression in Male 1 was significantly higher than in Male 3 (p = 0.0032, Male 1 = 0.8361, Male 3 = 0.3032).

We compared our samples (Saline-males, PGE2-males, Male 1, Male 2, Male 3) to one another for in β-actin expression levels (Fig 10C). Technical replicates were assigned as a random effect. Again, we found a significant reduction in β-actin expression levels in PGE2-exposed males compared to Saline male controls (p = 0.0020, Saline = 1.000, PGE2 = 0.3556). Compared to the Saline male controls pooled the expression of β-actin was reduced in Male 1 (p = 0.0206, Saline = 1.000, Male 1 = 0.5644), Male 2 (p = 0.0007, Saline = 1.000, Male 2 = 0.2661), and Male 3 (p = 0.0008, Saline = 1.000, Male 3 = 0.2749). However, we found no significant difference between PGE2-males pooled and Male 1 (p = 0.2256, PGE2 = 0.3556, Male 1 = 0.5644), Male 2 (p = 0.5939, PGE2 = 0.3556, Male 2 = 0.2661), or Male 3 (p = 0.6304, PGE2 = 0.3556, Male 3= 0.2749). We also found no significant differences in β-actin expression between Males 1 and 2 (p = 0.0929, Male 1 = 0.5644, Male 2 = 0.2661), Males 2 and 3 (p = 0.9578, Male 2 = 0.2661, Male 3 = 0.2749) or between Males 1 and 3 (p = 0.1019, Male 1 = 0.5644, Male 3 = 0.2749).

Lastly, we compared expression of N-Cadherin between the samples (Saline-males, PGE2-males, Male 1, Male 2, Male 3), assigning technical replicates as a random effect (Fig 10D). As expected, we found a significant reduction in N-Cadherin expression levels in PGE2-exposed males pooled compared to Saline male controls (p = 0.0203, Saline = 1.000, PGE2 = 0.5519). The expression of N-Cadherin in Male 1 was not significantly different from the Saline male controls pooled (p = 0.5504, Saline = 1.000, Male 1 = 1.1030) but unexpectedly higher than in the PGE2-male pool (p = 0.0065, PGE2 = 0.5519, Male 1 = 1.1030). The expression of N-cadherin in Male 3 was significantly lower than the Saline male controls pooled (p = 0.0001. Saline = 1.000, Male 2 = 0.0499), Male 1 (p < 0.001, Male 1 = 1.1030, Male 2 = 0.0499), but not significantly different from Male 3 (p = 0.4024, Male 2 = 0.0499, Male 3 = 0.1954). Similarly, the expression level in Male 3 was significantly lower than the Saline male controls pooled (p = 0.0001. Saline = 1.000, Male 3 = 0.1954) and Male 1 (p < 0.001, Male 1 = 1.1030, Male 2 = 0.0499). Surprisingly, compared to the PGE2-exposed males pooled, there was a reduction in N-Cadherin expression in Male 2 (p = 0.0111, PGE2 = 0.5519, Male 2 = 0.0499).

In summary, we showed that even though PGE2-exposed male mice on average had reduced expressions of spinophilin, β-actin, and N-Cadherin, the levels of these proteins vary depending on individual behavioural outcomes. The PGE2-exposed Male 1 which performed the best across cerebellar motor tests had expression levels of spinophilin and N-Cadherin similar to Saline control mice, while those that performed the worst across cerebellar motor tests (Males 2 and 3) had expression levels of spinophilin and N-Cadherin more similar to the PGE2-exposed mice.

## Discussion

Clinical and epidemiological studies have documented that numerous environmental risk factors affect the levels of PGE2 at various points of prenatal development and result in ASDs [10, 12, 108]. The molecular mechanisms by which changes in prenatal PGE2 levels affect neuronal development and lead to ASD symptoms postnatally are still largely unknown. In this study we demonstrated for the first time that a single maternal injection of PGE2 in mice during a critical time point in pregnancy affects the developing cerebellum in offspring, including cell density, actin-dependent morphology of dendrites and dendritic spines, the expression of cytoskeletal proteins and relevant behavioural outcomes postnatally. While literature provides some insight into the consequences of altered COX2/PGE2 signalling during early postnatal stages of the cerebellum development [54, 109], our unique approach investigates the effects of prenatal exposure to PGE2. Importantly, many of the molecular and behavioural changes observed here were sex dependent.

The cerebellum has been frequently implicated in the etiology of ASD, with prenatal and early postnatal disruption of neuronal circuits having a strong positive correlation to later development of symptoms [110–113]. Our findings provide evidence that exposure to PGE2 during critical time in pregnancy can disrupt normal cerebellar development in offspring, resulting in (1) reduced density of cells originating at G11 (in both males and females) and G16 (in females alone), followed by (2) increased dendritic arborization, looping, spine density and odds of observing mature spines, (3) decreased expression level of cytoskeletal proteins, and (4) impaired related motor coordination postnatally(Table 3). We discuss these deficits and sex differences below.

**Table 3:** Summary of PGE2 effects on cerebellar dendritic morphology, protein expression and behaviours. Dendritic morphology findings are summarized. The effect in PGE2-exposed male and females are shown compared to sex matched Saline controls. Sex differences within PGE2 and Saline mice are indicated by up and down arrows. Sex differences in PGE2-exposed animals that were different from the Saline controls are indicated in **bold**.

| Measure | PGE2 Male | PGE2 Female | Sex differences<br>M=Male; F=Female; ↑=higher; ↓=lower |
| --- | --- | --- | --- |
| **Cell Density** |  |  |  |
| G11 cells | ↓ | ↓ | Saline: M↓; F↑<br>PGE2: M↓; F↑ |
| G16 cells | - | ↓ | Saline: M↓; F↑<br>**PGE2: No sex-difference** |
| **Dendritic Morphology** |  |  |  |
| Dendrite Length | - | - | Saline: M↓; F↑<br>**PGE2: No sex-difference** |
| Dendritic Arborization | ↑ | ↑ | Saline and PGE2: No sex-difference |
| Dendritic Looping | ↑ | ↑ | Saline and PGE2: No sex-difference |
| Spine Density | ↑ | ↑ | Saline and PGE2: No sex-difference |
| Odds of Mature Spines | ↑ | ↑ | Saline: M↑; F↓<br>**PGE2: No sex-difference** |
| **Motor Behaviour** |  |  |  |
| Adhesive Sticker Test:<br>Swipes per Second | ↑ | - | Saline: No sex-difference<br>**PGE2: M↑; F↓** |
| Grid Walking Test:<br># of Slips | - | - | Saline: No sex-difference<br>**PGE2: M↑; F↓** |
| Grid Walking Test:<br>Time to Escape | - | - | Saline: M↓; F↑<br>**PGE2: M↑; F↓** |
| Cylinder Test:<br># of Forelimb Touches | - | - | Saline: M↓; F↑<br>**PGE2: No sex-difference** |
| Cylinder Test:<br># of Rears | - | - | Saline: M↓; F↑<br>**PGE2: No sex-difference** |
| Cylinder Test:<br># of Steps | ↓ | - | Saline and PGE2: No sex-difference |
| **Cytoskeletal Protein Expression** |  |  |  |
| Spinophilin | ↓ | ↓ | Saline: M↑; F↓<br>**PGE2: No sex-difference** |
| β-Actin | ↓ | ↓ | Saline: M↑; F↓<br>**PGE2: No sex-difference** |
| N-Cadherin | ↓ | ↓ | Saline and PGE2: No sex-difference |

### The effect of PGE2 on cell density and dendritic morphology in the cerebellum

In this study we observed that the maternal exposure to PGE2 during the critical time in development at G11, causes a reduction of cell density of two cohorts of cells originating at G11 and G16 in the cerebellum. Specifically, we found that PGE2-exposed males and females had a lower density of G11-labelled cells compared to sex-matched saline controls. PGE2-exposed females also exhibited a lower density of G16-labelled cerebellar cells than controls. One of the most consistently reported abnormalities of the brain in ASD cases is the decreased number of Purkinje cells in the cerebellum [114, 115] and reductions in Purkinje cell size [116–118]. Moreover, cerebellar dysplasia [119] and a reduction in total cerebellar volume in children [120] and adults [121] with ASD have also been reported. It is feasible that the reported cerebellar abnormalities observed in ASD cases may be attributed to diminished cell numbers due to altered cell proliferation in early development [49, 122].

Our previous *in vitro* study have demonstrated that PGE2 exposure of neuroectodermal (NE-4C) cells during differentiation increases dendrite length. [86]. We did not observe any difference in dendrite length in the cerebellum with PGE2-exposure. However, we observed that the innate sex-difference within saline controls with females having significantly longer dendrites than males was no longer present in PGE2-exposed animals (Table 3). Similar data comes from studies in rats showing that healthy females have longer dendritic length compared to males within certain areas of the brain including the locus coeruleus [123] and the hippocampus [124]. Thus, the lack of expected sex-differences in PGE2-exposed mice might suggest a potential influence of PGE2-exposure of dendrite length in male or female mice.

Although there was no apparent difference in dendrite length in PGE2-exposed mice, we observed an increase in dendritic arborization closer to the soma at 20-50µm compared to saline control mice. Sex-differences were not seen in either saline controls or PGE2-exposed mice. Other studies found that a local injection of PGE2 into the rat cerebellum during the second postnatal week stunted dendritic arborization [54, 94]. In contrast, the injection of COX-2 inhibitors into the rat cerebellum during the same time point increased dendritic arborization [54]. In addition, the observed changes in dendritic arbors and cerebellar synaptic density reported in PGE2-exposured rats also had increased estriol (E2) levels [54, 94] that subsequently resulted in stunted dendritic arbors and a reduction in cerebellar synaptic density [46]. The PGE2-E2 collaborative effect also resulted in increased dendritic arborization within the prefrontal cortex of the postnatally PGE2-injected rats [125, 126].

We have previously observed an increase in dendrite looping *in vitro* in differentiating NE-4C cells treated with PGE2 [86] and increased odds of finding looping dendrites in the cerebellums of COX-2^-^-KI mice [56]. In this study, we observed that PGE2-exposed male and female offspring also exhibit increased the odds of observing dendritic looping. The formation of dendritic loops is often associated with dysregulation of the polymerization rate or in the direction of polymerization of the actin cytoskeletal dynamics, leading to disruptions in self-avoidance mechanisms and resulting in abnormal dendritic pathfinding [127, 128]. The increase in dendritic loop formation is often a sign of disruptions in dendritic outgrowth and pathfinding which are linked to neurodevelopmental disorders including retinal dysplasia [129], and ASD [130].

With the increased dendritic arborization, we also observed increased dendritic spine density in both PGE2-exposed male and female mice compared to sex-matched saline controls with no apparent sex-differences (Table 3). Higher dendritic spine density typically corresponds to an increase in connectivity between neurons [131]. Abnormal Purkinje cell dendritic spine density was also observed in other autism-related mouse models, includingShank3+/ΔC, Mecp2^R308/Y^ [132] and the Valproic acid-injected [133] as well as in humans with fragile X syndrome [134–136]. Changes in dendritic spine density have been associated with excitatory/inhibitory (E/I) imbalances that are common in the brains of ASD individuals and in many genetic ASD *in vitro* and *in vivo* rodent models [137–139]. Interestingly, increases in excitatory signalling correspond to a decrease in dendritic spine density [140, 141]. The cooccurrence of E/I imbalances and changes in dendritic spine density are consistent in other mouse models of ASD [142–144], including in the Rett syndrome knockout mice (*Mecp2* KO), and *mTOR* KO mice [145]. This existing evidence is not surprising given the contribution of dendritic spines to the strength and plasticity of excitatory synapses [146], and suggest that prenatal PGE2-exposure might potentially create an E/I imbalance in the cerebellum of offspring mice.

In addition to the increased spine density and arborization PGE2-exposed mice also have increased odds of seeing mature spines. Normally dendritic spines can be found in different shapes which are related to their function, and are categorized as thin, stubby (less mature) and mushroom (mature) shaped [97, 147, 148]. We reported dendritic morphology by examining the likelihood of observing a mature spine (mushroom) compared to an immature spine (thin or stubby). First, we observed that in saline controls there was a decrease in the odds of observing mature (mushroom shaped) spines in females compared to males. This decrease was not unexpected given that fluctuations in dendritic spine morphology (between mature and immature) have been reported in adult female rats due to the estrus cycle [149, 150]. Being the intermediary shape between thin and mushroom shaped spines, the increase in stubby shaped spines in Saline control females compared to males may speak to the increase in shifts between thin and mushroom spines. PGE2-exposed males and females were both more likely to have more mature (mushroom shaped) spines compared to Saline controls with sex-differences no longer present. The loss of the sex difference indicates that there may be a reduction in estrus-dependent changes in spine shape in PGE2-exposed females. Studies in postnatal rats showed that an increase in E2 increased the relative quantities of mushroom shaped spines in CA1 in the hippocampus [151]. Building on previous findings that PGE2-exposure can increase E2 levels, we speculate that prenatal exposure to environmental risk factors known to affect PGE2 levels may contribute to brain pathologies in early development via impacting E2 levels. Given the role of E2 in brain masculinization during neurodevelopment both prenatally, and postnatally, these findings are in line with the inherent sex-bias in ASD-this is redundant to what was said above

### Abnormal motor behaviour in PGE2-exposed offspring

Previous studies [in humans?] showed that early disruption of cerebellar neural circuitry is associated with diagnosis of ASD [152–154]. As discussed above, rodent models have also shown that disruptions in cerebellar morphology including increases or decreases in cell density and size, dendritic arborization, dendritic spine density, and changes in dendritic spine morphology results in ASD related behaviours including social deficits, difficulties in task-switching and anxious grooming behaviours [155–157], with some studies showing that greater disruptions cerebellar morphology corresponded to greater severity of ASD-related behaviours [112, 158]. We previously demonstrated that PGE2-exposed mice offspring also exhibit autism-related behaviours [38, 39]. To build on those findings we further investigated whether the PGE2-exposed mice offspring also exhibit specific cerebellar-related motor dysfunction. We observed a male specific effect on sensorimotor coordination in the adhesive sticker test and a reduction in locomotion in the cylinder test. In addition, we found clear sex differences within PGE2-exposed animals which were normally not present in the controls.

In the *adhesive sticker test*, which is a sensitive test for sensorimotor dysfunction [78, 159, 160] we found a male-specific increase in the number of swipes per second, indicating that PGE2-exposed males had difficulties in sensory-motor integration. Basic motor dysfunction in ASD is often attributed to deficits in sensorimotor integration in which individuals are unable to extract and integrate sensory information into an executable motor plan [161–163]. Literature suggests that these deficits in motor coordination arise from disruptions in cortico-cerebellar networks [161, 164]. Our findings suggest that PGE2 exposure may make males more inefficient in the execution of motor planning, resulting in an excess of swipes before the adhesive sticker is removed. The *grid walking test* (or foot-fault test) is an established measure of sensorimotor coordination of the four limbs in rodent models [81, 82]. In the number of slips (number-of-faults) and the time to escape we show that PGE2-exposure may affect sensorimotor coordination differently in males and females with males slipping more but escaping more quickly than females. Whereas the saline controls show no sex difference in the number of slips but males seem to escape more slowly than females. The *adhesive sticker* and the *grid waling tests* together suggest that cerebellar motor performance may be more disrupted in PGE2-exposed males than females.

We also conducted the *cylinder test* to measure cerebellar-dependent spontaneous forelimb and hindlimb use as well as the postural control necessary for rearing in mice [83–85]. Within saline controls males made less forelimb touches and reared less frequently than females (Table 3). This is consistent with other studies showing that male mice inherently rear less than females [165]. However, the expected sex differences in forelimb touches and rearing were no longer observed in PGE2-exposed mice. In addition, we observed a male specific decrease in number of steps in PGE2-exposed mice compared to saline controls with no sex differences were observed in either condition. Deficits in cerebellar-related motor locomotion have been reported in ASD individuals [166–169], however these behavioural impairments are not well researched in rodent models of autism.

### Abnormal levels of essential cytoskeletal proteins

We have previously demonstrated that abnormalities of neurite and neurite morphology in PGE2-exposed differentiating NE-4C stem cells *in vitro* [86] and COX-2^-^-KI mice [56] are associated with abnormal expression levels of β-actin (decreased in COX-2^-^-KI mice) and actin-binding protein spinophilin (increased in NE-4C stem cells and COX-2^-^-KI mice). In this study, PGE2-exposed mice had decreased expression levels of β-actin, spinophilin, and a cell adhesion molecule called N-cadherin in both males and females relative to sex-matched controls. Within both saline control mice, we observed a higher expression of both β-actin and spinophilin in males compared to females, which was not present in PGE2-exposed mice. We did not observe any sex-differences in N-Cadherins in PGE2-exposed or Saline control mice.

β-actin and spinophilin play key roles in the maintenance of the actin-cytoskeleton. The rate of actin-polymerisation/depolymerization affects the shape and growth of dendritic spines and is a determinant of dendrite pathfinding [170–174]. The primary function of spinophilin is in the stabilization of the actin cytoskeleton and is involved in actin driven changes in dendrite and dendritic spine morphology as result of excitatory synaptic activity [175, 176]. Here we observed an increase in density of more mature spines (or less immature spines) in PGE2-exposed mice that coincided with a reduction in the expression level of β-actin and spinophilin. This is in line with a previous study in rats injected with PGE2 postnatally which found decreased spinophilin protein in the cerebellums [94]. Another study also reported an increase in dendritic spine density in spinophilin-knockout mice [177]. The reduction in β-actin and spinophilin protein content coupled with the increased odds of dendrite looping might indicate cytoskeletal dysfunction within the cerebellums of PGE2-exposed mice, which may subsequently contribute to the abnormal cerebellar motor behaviour observed in the offspring.

Lastly, PGE2-exposed male and female mice also had a reduction in the expression levels of N-Cadherin, a well-known cell adhesion molecule with important roles throughout neural development [105, 178–180]. N-Cadherins have also known roles in dendritic outgrowth and guidance, and synaptogenesis and synaptic plasticity [106, 107]. Mutations in N-Cadherin including copy number variations and single nucleotide polymorphisms are associated with ASD [181, 182]. Interestingly, previous findings in cultured neurons showed that an inhibition of N-Cadherin by use of a dominant-negative N-Cadherin (cN390Δ), resulted in a filopodia-like elongation of dendritic spines, and a disruption of the distribution of proteins on the postsynaptic membrane [103, 104]. We found no sex-differences between either Saline control, or PGE2-exposed males or females. We also observed a correlation between the expression levels of these three proteins and behavioural outcomes in PGE2-exposed males, found to be the most affected group in cerebellar-motor function. We determined that the expression levels of spinophilin and N-cadherin in Male 1 (which had the best overall performance among PGE2 males) was close to the average seen in the saline control males whereas Males 2 and 3 (worst scoring PGE2-exposed males) had expression levels similar to PGE2-exposed males.

Our previous studies in the same mouse model already demonstrated that maternal exposure to PGE2 resulted in a cross-talk between the COX2/PGE2 signalling and canonical Wnt signalling pathways, causing differential expression of ASD and Wnt genes in genetically identical offspring [87] and manifestation of sex-dependent autism behaviour [38]. Our current findings provide further evidence that the male and female offspring prenatally exposed to PGE2 also exhibit abnormal dendritic morphology within the cerebellum associated with deficits in motor behaviour.

### Conclusion remarks

To summarize, we show for the first time that a one-time maternal exposure to PGE2 results in disruptions in cell density, and dendritic and dendritic spine morphology postnatally in males and females offspring. These morphological changes corresponded to abnormal expression levels of β-actin, spinophilin, and N-cadherin Additionally, PGE2-exposed mice show deficits on cerebellar motor function tests. Examining PGE2 mice we found that variability of performance on cerebellar motor function tests corresponded to the expression of spinophilin and N-cadherin. These findings build on previous clinical examples that maternal exposure to common environmental factors that affect PGE2 levels during critical time points have consequences for the developing brain and are manifested in spectrum of postnatal behaviour. The most challenging aspect of studying the molecular basis of neurodevelopmental disorders is finding a suitable model system. Our studies in PGE2-exposed and COX2-KI mice along with research in other model systems described above add to the growing evidence that abnormal COX2/PGE2 signaling during critical time points in development may be one of the contributing causes of ASD. This study also emphasizes the importance of assessing sex-differences especially relevant to disorders such as ASD, which are still greatly underrepresented in the literature.

## Acknowledgements

This research was funded by the Natural Sciences and Engineering Research Council of Canada (NSERC).

## Author Contributions

AK collected and analyzed the data, created figures and tables, and wrote the manuscript. KH performed behavioural testing of mice. CW stained, imaged and quantified cell density in G11 and G16 cerebellar slides. DAC supervised the design and coordination of the study and was involved with writing the manuscript.

## References

1. Tassoni, D., et al., The role of eicosanoids in the brain. Asia Pac J Clin Nutr, 2008. 17 **Suppl 1**: p. 220–8.

2. Chen, C. and N.G. Bazan, Lipid signaling: sleep, synaptic plasticity, and neuroprotection. Prostaglandins Other Lipid Mediat, 2005. 77(1-4): p. 65–76.

3. Sang, N. and C. Chen, Lipid signaling and synaptic plasticity. Neuroscientist, 2006. 12(5): p. 425–34.

4. Park, J.Y., M.H. Pillinger, and S.B. Abramson, Prostaglandin E2 synthesis and secretion: the role of PGE2 synthases. Clin Immunol, 2006. 119(3): p. 229–40.

5. Rouzer, C.A. and L.J. Marnett, Cyclooxygenases: structural and functional insights. Journal of lipid research, 2009. 50(Supplement): p. S29–S34.

6. Hoozemans, J.J., et al., Cyclooxygenase expression in microglia and neurons in Alzheimer’s disease and control brain. Acta Neuropathol, 2001. 101(1): p. 2–8.

7. Schwab, J.M., et al., Selective accumulation of cyclooxygenase-1-expressing microglial cells/macrophages in lesions of human focal cerebral ischemia. Acta Neuropathol, 2000. 99(6): p. 609–14.

8. Kirkby, N.S., et al., Systematic study of constitutive cyclooxygenase-2 expression: Role of NF-kappaB and NFAT transcriptional pathways. Proc Natl Acad Sci U S A, 2016. 113(2): p. 434–9.

9. Maslinska, D., et al., Constitutive expression of cyclooxygenase-2 (COX-2) in developing brain. A. Choroid plexus in human fetuses. Folia Neuropathol, 1999. 37(4): p. 287–91.

10. Tamiji, J. and D.A. Crawford, The neurobiology of lipid metabolism in autism spectrum disorders. Neurosignals, 2010. 18(2): p. 98–112.

11. Wong, C. and D.A. Crawford, Lipid signalling in the pathology of autism spectrum disorders. Comprehensive guide to autism, 2014. 18: p. 1259–1283.

12. Wong, C.T., J. Wais, and D.A. Crawford, Prenatal exposure to common environmental factors affects brain lipids and increases risk of developing autism spectrum disorders. European Journal of Neuroscience, 2015. 42(10): p. 2742–2760.

13. Yoo, H.J., et al., Association between PTGS2 polymorphism and autism spectrum disorders in Korean trios. Neurosci Res, 2008. 62(1): p. 66–9.

14. Volk, H.E., et al., Residential proximity to freeways and autism in the CHARGE study. Environ Health Perspect, 2011. 119(6): p. 873–7.

15. Costa, L.G., et al., Neurotoxicity of traffic-related air pollution. Neurotoxicology, 2017. 59: p. 133–139.

16. Samsel, A. and S. Seneff, Glyphosate, pathways to modern diseases III: Manganese, neurological diseases, and associated pathologies. Surg Neurol Int, 2015. 6: p. 45.

17. Suzuki, K., et al., Plasma cytokine profiles in subjects with high-functioning autism spectrum disorders. PLoS One, 2011. 6(5): p. e20470.

18. Larsson, M., et al., Associations between indoor environmental factors and parental-reported autistic spectrum disorders in children 6-8 years of age. Neurotoxicology, 2009. 30(5): p. 822–31.

19. Miodovnik, A., et al., Endocrine disruptors and childhood social impairment. Neurotoxicology, 2011. 32(2): p. 261–7.

20. Testa, C., et al., Di-(2-ethylhexyl) phthalate and autism spectrum disorders. ASN Neuro, 2012. 4(4): p. 223–9.

21. Andrade, C., Use of acetaminophen (paracetamol) during pregnancy and the risk of autism spectrum disorder in the offspring. J Clin Psychiatry, 2016. 77(2): p. e152–4.

22. Avella-Garcia, C.B., et al., Acetaminophen use in pregnancy and neurodevelopment: attention function and autism spectrum symptoms. Int J Epidemiol, 2016. 45(6): p. 1987–1996.

23. Bauer, A.Z., et al., Prenatal paracetamol exposure and child neurodevelopment: A review. Horm Behav, 2018. 101: p. 125–147.

24. Bittker, S.S. and K.R. Bell, Acetaminophen, antibiotics, ear infection, breastfeeding, vitamin D drops, and autism: an epidemiological study. Neuropsychiatr Dis Treat, 2018. 14: p. 1399–1414.

25. Bittker, S.S. and K.R. Bell, Postnatal Acetaminophen and Potential Risk of Autism Spectrum Disorder among Males. Behav Sci (Basel), 2020. 10(1).

26. Brandlistuen, R.E., et al., Prenatal paracetamol exposure and child neurodevelopment: a sibling-controlled cohort study. Int J Epidemiol, 2013. 42(6): p. 1702–13.

27. Liew, Z., et al., Maternal use of acetaminophen during pregnancy and risk of autism spectrum disorders in childhood: A Danish national birth cohort study. Autism Res, 2016. 9(9): p. 951–8.

28. Schultz, S.T. and G.G. Gould, Acetaminophen Use for Fever in Children Associated with Autism Spectrum Disorder. Autism Open Access, 2016. 6(2).

29. Alemany, S., et al., Prenatal and postnatal exposure to acetaminophen in relation to autism spectrum and attention-deficit and hyperactivity symptoms in childhood: Meta-analysis in six European population-based cohorts. Eur J Epidemiol, 2021.

30. Bandim, J.M., et al., Autism and Möbius sequence: an exploratory study of children in northeastern Brazil. Arq Neuropsiquiatr, 2003. 61(2a): p. 181–5.

31. Bos-Thompson, M.A., et al., Möbius syndrome in a neonate after mifepristone and misoprostol elective abortion failure. Ann Pharmacother, 2008. 42(6): p. 888–92.

32. Wong, C.T., J. Wais, and D.A. Crawford, Prenatal exposure to common environmental factors affects brain lipids and increases risk of developing autism spectrum disorders. Eur J Neurosci, 2015. 42(10): p. 2742–60.

33. Tamiji, J. and D.A. Crawford, Prostaglandin E2 and misoprostol induce neurite retraction in Neuro-2a cells. Biochemical and biophysical research communications, 2010. 398(3): p. 450–456.

34. Davidson, J.M., et al., Prostaglandin E2 elevates calcium in differentiated neuroectodermal stem cells. Molecular and Cellular Neuroscience, 2016. 74: p. 71–77.

35. Wong, C.T., et al., Prostaglandin E2 alters Wnt-dependent migration and proliferation in neuroectodermal stem cells: implications for autism spectrum disorders. Cell Commun Signal, 2014. 12: p. 19.

36. Wong, C.T., et al., Prostaglandin E2 promotes neural proliferation and differentiation and regulates Wnt target gene expression. J Neurosci Res, 2016. 94(8): p. 759–75.

37. Wong, C.T., I. Bestard-Lorigados, and D.A. Crawford, Autism-related behaviors in the cyclooxygenase-2-deficient mouse model. Genes Brain Behav, 2019. 18(1): p. e12506.

38. Wong, C.T., et al., *Abnormal prostaglandin E2 signalling results in autism-associated behaviours in novel mouse models*, in *Society for Neuroscience* 2017: Washington D.C.

39. Kissoondoyal, A., et al., Unpublished.

40. Rai-Bhogal, R., et al., Maternal exposure to prostaglandin E2 modifies expression of Wnt genes in mouse brain - An autism connection. Biochem Biophys Rep, 2018. 14: p. 43–53.

41. Battle, D.E., Diagnostic and Statistical Manual of Mental Disorders (DSM). Codas, 2013. 25(2): p. 191–2.

42. Lai, M.C., et al., Sex/gender differences and autism: setting the scene for future research. J Am Acad Child Adolesc Psychiatry, 2015. 54(1): p. 11–24.

43. Teitelbaum, P., et al., Movement analysis in infancy may be useful for early diagnosis of autism. Proceedings of the National Academy of Sciences, 1998. 95(23): p. 13982–13987.

44. Minshew, N.J., et al., Underdevelopment of the postural control system in autism. Neurology, 2004. 63(11): p. 2056–2061.

45. Kohen-Raz, R., F.R. Volkman, and D.J. Cohen, Postural control in children with autism. Journal of autism and developmental disorders, 1992. 22(3): p. 419–432.

46. Ghaziuddin, M. and E. Butler, Clumsiness in autism and Asperger syndrome: A further report. Journal of Intellectual Disability Research, 1998. 42(1): p. 43–48.

47. Miyahara, M., et al., Brief report: motor incoordination in children with Asperger syndrome and learning disabilities. Journal of autism and developmental disorders, 1997. 27(5): p. 595–603.

48. Mari, M., et al., The reach–to–grasp movement in children with autism spectrum disorder. Philosophical Transactions of the Royal Society of London. Series B: Biological Sciences, 2003. 358(1430): p. 393–403.

49. Courchesne, E., et al., The ASD Living Biology: from cell proliferation to clinical phenotype. Mol Psychiatry, 2019. 24(1): p. 88–107.

50. Beversdorf, D.Q., et al., Timing of prenatal stressors and autism. J Autism Dev Disord, 2005. 35(4): p. 471–8.

51. Courchesne, E., et al., Unusual brain growth patterns in early life in patients with autistic disorder: an MRI study. Neurology, 2001. 57(2): p. 245–54.

52. Hashimoto, T., et al., Development of the brainstem and cerebellum in autistic patients. J Autism Dev Disord, 1995. 25(1): p. 1–18.

53. Limperopoulos, C., et al., Does cerebellar injury in premature infants contribute to the high prevalence of long-term cognitive, learning, and behavioral disability in survivors? Pediatrics, 2007. 120(3): p. 584–93.

54. Dean, S.L., et al., Prostaglandin E2 is an endogenous modulator of cerebellar development and complex behavior during a sensitive postnatal period. Eur J Neurosci, 2012. 35(8): p. 1218–29.

55. Hoffman, J.F., C.L. Wright, and M.M. McCarthy, A Critical Period in Purkinje Cell Development Is Mediated by Local Estradiol Synthesis, Disrupted by Inflammation, and Has Enduring Consequences Only for Males. J Neurosci, 2016. 36(39): p. 10039–49.

56. Kissoondoyal, A., R. Rai-Bhogal, and D.A. Crawford, Abnormal dendritic morphology in the cerebellum of cyclooxygenase-2-knockin mice. Eur J Neurosci, 2021. 54(7): p. 6355–6373.

57. Ohno, T., H. Ohtsuki, and S. Okabe, Effects of 16,16-dimethyl prostaglandin E2 on ethanol-induced and aspirin-induced gastric damage in the rat. Scanning electron microscopic study. Gastroenterology, 1985. 88(1 Pt 2): p. 353–61.

58. Steffenrud, S., *Metabolism of 16*, *16-dimethyl-prostaglandin E2 in the human female*. Biochem Med, 1980. 24(3): p. 274–92.

59. Hagenbuch, B. and P.J. Meier, Organic anion transporting polypeptides of the OATP/ SLC21 family: phylogenetic classification as OATP/ SLCO superfamily, new nomenclature and molecular/functional properties. Pflugers Arch, 2004. 447(5): p. 653–65.

60. Roth, M., A. Obaidat, and B. Hagenbuch, *OATPs,* OATs and OCTs: the organic anion and cation transporters of the SLCO and SLC22A gene superfamilies. Br J Pharmacol, 2012. 165(5): p. 1260–87.

61. Löscher, W. and H. Potschka, Role of drug efflux transporters in the brain for drug disposition and treatment of brain diseases. Prog Neurobiol, 2005. 76(1): p. 22–76.

62. Semple, B.D., et al., Brain development in rodents and humans: Identifying benchmarks of maturation and vulnerability to injury across species. Prog Neurobiol, 2013. 106**-****107**: p. 1–16.

63. Takahashi, T., R.S. Nowakowski, and V.S. Caviness, Jr., The cell cycle of the pseudostratified ventricular epithelium of the embryonic murine cerebral wall. J Neurosci, 1995. 15(9): p. 6046–57.

64. Rodier, P.M., Chronology of neuron development: animal studies and their clinical implications. Dev Med Child Neurol, 1980. 22(4): p. 525–45.

65. Clancy, B., R.B. Darlington, and B.L. Finlay, Translating developmental time across mammalian species. Neuroscience, 2001. 105(1): p. 7–17.

66. Clancy, B., et al., Extrapolating brain development from experimental species to humans. Neurotoxicology, 2007. 28(5): p. 931–7.

67. Pastuszak, A.L., et al., Use of misoprostol during pregnancy and Möbius’ syndrome in infants. N Engl J Med, 1998. 338(26): p. 1881–5.

68. Tuttle, A.H., et al., Immunofluorescent detection of two thymidine analogues (CldU and IdU) in primary tissue. Journal of visualized experiments : JoVE, 2010(46): p. 2166.

69. Tuttle, A.H., et al., Immunofluorescent detection of two thymidine analogues (CldU and IdU) in primary tissue. J Vis Exp, 2010(46).

70. Zaqout, S. and A.M. Kaindl, Golgi-Cox Staining Step by Step. Front Neuroanat, 2016. 10: p. 38.

71. Longair, M.H., D.A. Baker, and J.D. Armstrong, Simple Neurite Tracer: open source software for reconstruction, visualization and analysis of neuronal processes. Bioinformatics, 2011. 27(17): p. 2453–2454.

72. Schindelin, J., et al., Fiji: an open-source platform for biological-image analysis. Nature Methods, 2012. 9(7): p. 676–682.

73. Rueden, C.T., et al., ImageJ2: ImageJ for the next generation of scientific image data. BMC Bioinformatics, 2017. 18(1): p. 529.

74. Kissoondoyal, A., R. Rai-Bhogal, and D.A. Crawford, A Quantification of Dendritic and dendritic spine morphology in Cyclooxygenase - 2 – Knockin Mice, in Neuromatch. 2020: Virtual.

75. Bae, J., et al., NESH regulates dendritic spine morphology and synapse formation. PLoS One, 2012. 7(4): p. e34677.

76. Briones, B.A., et al., Response learning stimulates dendritic spine growth on dorsal striatal medium spiny neurons. Neurobiol Learn Mem, 2018. 155: p. 50–59.

77. Sorge, R.E., et al., Olfactory exposure to males, including men, causes stress and related analgesia in rodents. Nat Methods, 2014. 11(6): p. 629–32.

78. Richter, F., et al., Sensorimotor tests unmask a phenotype in the DYT1 knock-in mouse model of dystonia. Behav Brain Res, 2017. 317: p. 536–541.

79. Lam, H.A., et al., Elevated tonic extracellular dopamine concentration and altered dopamine modulation of synaptic activity precede dopamine loss in the striatum of mice overexpressing human α-synuclein. J Neurosci Res, 2011. 89(7): p. 1091–102.

80. Fleming, S.M., et al., Early and progressive sensorimotor anomalies in mice overexpressing wild-type human alpha-synuclein. J Neurosci, 2004. 24(42): p. 9434–40.

81. Heyser, C.J., Assessment of developmental milestones in rodents. Curr Protoc Neurosci, 2004. **Chapter 8**: p. Unit 8.18.

82. Feather-Schussler, D.N. and T.S. Ferguson, A Battery of Motor Tests in a Neonatal Mouse Model of Cerebral Palsy. J Vis Exp, 2016(117).

83. Schönfeld, L.M., et al., Long-Term Motor Deficits after Controlled Cortical Impact in Rats Can Be Detected by Fine Motor Skill Tests but Not by Automated Gait Analysis. J Neurotrauma, 2017. 34(2): p. 505–516.

84. Schönfeld, L.M., et al., Evaluating rodent motor functions: Which tests to choose? Neurosci Biobehav Rev, 2017. 83: p. 298–312.

85. Rattka, M., et al., A Novel Approach to Assess Motor Outcome of Deep Brain Stimulation Effects in the Hemiparkinsonian Rat: Staircase and Cylinder Test. J Vis Exp, 2016(111).

86. Kissoondoyal, A. and D.A. Crawford, Prostaglandin E2 Increases Neurite Length and the Formation of Axonal Loops, and Regulates Cone Turning in Differentiating NE4C Cells Via PKA. Cellular and Molecular Neurobiology, 2021.

87. Rai-Bhogal, R., et al., Maternal exposure to prostaglandin E(2) modifies expression of Wnt genes in mouse brain - An autism connection. Biochem Biophys Rep, 2018. 14: p. 43–53.

88. Team, R.C., R: A language and environment for statistical computing. 2013, R Foundation for Statistical Computing: Vienna, Austria.

89. Croissant, Y., Estimation of Random Utility Models in R: The mlogit Package. 2020, 2020. 95(11): p. 41.

90. Bates, D., et al., Fitting Linear Mixed-Effects Models Using lme4. 2015, 2015. 67(1): p. 48.

91. Faul, F., et al., G*Power 3: a flexible statistical power analysis program for the social, behavioral, and biomedical sciences. Behav Res Methods, 2007. 39(2): p. 175–91.

92. Bird, A.D. and H. Cuntz, Dissecting Sholl Analysis into Its Functional Components. Cell Rep, 2019. 27(10): p. 3081–3096.e5.

93. Binley, K.E., et al., Sholl analysis: a quantitative comparison of semi-automated methods. J Neurosci Methods, 2014. 225: p. 65–70.

94. Dean, S.L., et al., Prostaglandin E2 stimulates estradiol synthesis in the cerebellum postnatally with associated effects on Purkinje neuron dendritic arbor and electrophysiological properties. Endocrinology, 2012. 153(11): p. 5415–27.

95. Lai, K.O. and N.Y. Ip, Structural plasticity of dendritic spines: the underlying mechanisms and its dysregulation in brain disorders. Biochim Biophys Acta, 2013. 1832(12): p. 2257–63.

96. Tashiro, A. and R. Yuste, Structure and molecular organization of dendritic spines. Histol Histopathol, 2003. 18(2): p. 617–34.

97. Bourne, J. and K.M. Harris, Do thin spines learn to be mushroom spines that remember? Curr Opin Neurobiol, 2007. 17(3): p. 381–6.

98. Sardi, S., et al., Dendritic Learning as a Paradigm Shift in Brain Learning. ACS Chem Neurosci, 2018. 9(6): p. 1230–1232.

99. Mel, B.W., J. Schiller, and P. Poirazi, Synaptic plasticity in dendrites: complications and coping strategies. Curr Opin Neurobiol, 2017. 43: p. 177–186.

100. Joensuu, M., V. Lanoue, and P. Hotulainen, Dendritic spine actin cytoskeleton in autism spectrum disorder. Prog Neuropsychopharmacol Biol Psychiatry, 2018. 84(Pt B): p. 362–381.

101. Penzes, P. and I. Rafalovich, Regulation of the actin cytoskeleton in dendritic spines. Synaptic Plasticity, 2012: p. 81–95.

102. Dent, E.W., S.L. Gupton, and F.B. Gertler, The growth cone cytoskeleton in axon outgrowth and guidance. Cold Spring Harb Perspect Biol, 2011. 3(3).

103. Togashi, H., et al., Cadherin regulates dendritic spine morphogenesis. Neuron, 2002. 35(1): p. 77–89.

104. Mendez, P., et al., N-cadherin mediates plasticity-induced long-term spine stabilization. J Cell Biol, 2010. 189(3): p. 589–600.

105. Suzuki, S.C. and M. Takeichi, Cadherins in neuronal morphogenesis and function. Dev Growth Differ, 2008. 50 **Suppl 1**: p. S119–30.

106. Hirano, S. and M. Takeichi, Cadherins in brain morphogenesis and wiring. Physiol Rev, 2012. 92(2): p. 597–634.

107. Takeichi, M. and K. Abe, Synaptic contact dynamics controlled by cadherin and catenins. Trends Cell Biol, 2005. 15(4): p. 216–21.

108. Wong and D.A. Crawford, Lipid Signalling in the Pathology of Autism Spectrum Disorders, in Comprehensive Guide to Autism, P. V., P. V., and M. C., Editors. 2014, Springer: New York, NY.

109. Wright, C.L., J.H. Hoffman, and M.M. McCarthy, Evidence that inflammation promotes estradiol synthesis in human cerebellum during early childhood. Transl Psychiatry, 2019. 9(1): p. 58.

110. Courchesne, E., et al., Autism at the beginning: microstructural and growth abnormalities underlying the cognitive and behavioral phenotype of autism. Dev Psychopathol, 2005. 17(3): p. 577–97.

111. Palmen, S.J., et al., Neuropathological findings in autism. Brain, 2004. 127(Pt 12): p. 2572–83.

112. D’Mello, A.M., et al., Cerebellar gray matter and lobular volumes correlate with core autism symptoms. Neuroimage Clin, 2015. 7: p. 631–9.

113. Mosconi, M.W., et al., The role of cerebellar circuitry alterations in the pathophysiology of autism spectrum disorders. Front Neurosci, 2015. 9: p. 296.

114. !!! INVALID CITATION !!! [50, 104].

115. Greco, C.M., et al., Neuropathologic features in the hippocampus and cerebellum of three older men with fragile X syndrome. Molecular autism, 2011. 2(1): p. 2–2.

116. Wegiel, J., et al., Contribution of olivofloccular circuitry developmental defects to atypical gaze in autism. Brain Res, 2013. 1512: p. 106–22.

117. Fatemi, S.H., et al., Purkinje cell size is reduced in cerebellum of patients with autism. Cell Mol Neurobiol, 2002. 22(2): p. 171–5.

118. Whitney, E.R., et al., Cerebellar Purkinje cells are reduced in a subpopulation of autistic brains: a stereological experiment using calbindin-D28k. Cerebellum, 2008. 7(3): p. 406–16.

119. Wegiel, J., et al., The neuropathology of autism: defects of neurogenesis and neuronal migration, and dysplastic changes. Acta Neuropathol, 2010. 119(6): p. 755–70.

120. Carper, R.A. and E. Courchesne, Inverse correlation between frontal lobe and cerebellum sizes in children with autism. Brain, 2000. 123(4): p. 836–844.

121. Hallahan, B., et al., Brain morphometry volume in autistic spectrum disorder: a magnetic resonance imaging study of adults. Psychol Med, 2009. 39(2): p. 337–46.

122. Marchetto, M.C., et al., Altered proliferation and networks in neural cells derived from idiopathic autistic individuals. Mol Psychiatry, 2017. 22(6): p. 820–835.

123. Bangasser, D.A., et al., Sexual dimorphism in locus coeruleus dendritic morphology: a structural basis for sex differences in emotional arousal. Physiol Behav, 2011. 103(3-4): p. 342–51.

124. McEwen, B.S., et al., Prevention of stress-induced morphological and cognitive consequences. Eur Neuropsychopharmacol, 1997. 7 **Suppl 3**: p. S323–8.

125. Garrett, J.E. and C.L. Wellman, Chronic stress effects on dendritic morphology in medial prefrontal cortex: sex differences and estrogen dependence. Neuroscience, 2009. 162(1): p. 195–207.

126. Radley, J.J., et al., Chronic behavioral stress induces apical dendritic reorganization in pyramidal neurons of the medial prefrontal cortex. Neuroscience, 2004. 125(1): p. 1–6.

127. Sdrulla, A.D. and D.J. Linden, Dynamic imaging of cerebellar Purkinje cells reveals a population of filopodia which cross-link dendrites during early postnatal development. Cerebellum, 2006. 5(2): p. 105–15.

128. Amthor, F.R. and C.W. Oyster, Spatial organization of retinal information about the direction of image motion. Proc Natl Acad Sci U S A, 1995. 92(9): p. 4002–5.

129. Weiner, J.A., et al., Axon fasciculation defects and retinal dysplasias in mice lacking the immunoglobulin superfamily adhesion molecule BEN/ALCAM/SC1. Mol Cell Neurosci, 2004. 27(1): p. 59–69.

130. Minshew, N.J. and D.L. Williams, The new neurobiology of autism: cortex, connectivity, and neuronal organization. Arch Neurol, 2007. 64(7): p. 945–50.

131. Fiala, J.C., J. Spacek, and K.M. Harris, Dendritic spine pathology: cause or consequence of neurological disorders? Brain research reviews, 2002. 39(1): p. 29–54.

132. Kloth, A.D., et al., Cerebellar associative sensory learning defects in five mouse autism models. Elife, 2015. 4: p. e06085.

133. Mychasiuk, R., et al., Effects of rat prenatal exposure to valproic acid on behaviour and neuro-anatomy. Dev Neurosci, 2012. 34(2-3): p. 268–76.

134. Irwin, S.A., et al., Abnormal dendritic spine characteristics in the temporal and visual cortices of patients with fragile-X syndrome: a quantitative examination. American journal of medical genetics, 2001. 98(2): p. 161–167.

135. Wisniewski, K., et al., The Fra (X) syndrome: neurological, electrophysiological, and neuropathological abnormalities. American journal of medical genetics, 1991. 38(2-3): p. 476–480.

136. Hinton, V., et al., Analysis of neocortex in three males with the fragile X syndrome. American journal of medical genetics, 1991. 41(3): p. 289–294.

137. Han, S., et al., Enhancement of inhibitory neurotransmission by GABAA receptors having α2,3-subunits ameliorates behavioral deficits in a mouse model of autism. Neuron, 2014. 81(6): p. 1282–1289.

138. Rubenstein, J.L. and M.M. Merzenich, Model of autism: increased ratio of excitation/inhibition in key neural systems. Genes Brain Behav, 2003. 2(5): p. 255–67.

139. Lee, E., J. Lee, and E. Kim, Excitation/Inhibition Imbalance in Animal Models of Autism Spectrum Disorders. Biol Psychiatry, 2017. 81(10): p. 838–847.

140. Gao, R. and P. Penzes, Common mechanisms of excitatory and inhibitory imbalance in schizophrenia and autism spectrum disorders. Curr Mol Med, 2015. 15(2): p. 146–67.

141. Takahashi, N., et al., Subcellular Imbalances in Synaptic Activity. Cell Rep, 2016. 14(6): p. 1348–1354.

142. Belichenko, P.V., et al., Widespread changes in dendritic and axonal morphology in Mecp2-mutant mouse models of Rett syndrome: evidence for disruption of neuronal networks. J Comp Neurol, 2009. 514(3): p. 240–58.

143. Tropea, D., et al., Partial reversal of Rett Syndrome-like symptoms in MeCP2 mutant mice. Proc Natl Acad Sci U S A, 2009. 106(6): p. 2029–34.

144. Chao, H.T., H.Y. Zoghbi, and C. Rosenmund, MeCP2 controls excitatory synaptic strength by regulating glutamatergic synapse number. Neuron, 2007. 56(1): p. 58–65.

145. Tang, G., et al., Loss of mTOR-dependent macroautophagy causes autistic-like synaptic pruning deficits. Neuron, 2014. 83(5): p. 1131–43.

146. Luebke, J.I., et al., Dendritic vulnerability in neurodegenerative disease: insights from analyses of cortical pyramidal neurons in transgenic mouse models. Brain Struct Funct, 2010. 214(2-3): p. 181–99.

147. Bourne, J.N. and K.M. Harris, Balancing structure and function at hippocampal dendritic spines. Annu Rev Neurosci, 2008. 31: p. 47–67.

148. Kasai, H., et al., Structure-stability-function relationships of dendritic spines. Trends Neurosci, 2003. 26(7): p. 360–8.

149. González-Burgos, I., M. Alejandre-Gómez, and M. Cervantes, Spine-type densities of hippocampal CA1 neurons vary in proestrus and estrus rats. Neuroscience letters, 2005. 379(1): p. 52–54.

150. de Lacalle, S., Estrogen effects on neuronal morphology. Endocrine, 2006. 29(2): p. 185–90.

151. Avila, J.A., et al., Estradiol rapidly increases GluA2-mushroom spines and decreases GluA2-filopodia spines in hippocampus CA1. Hippocampus, 2017. 27(12): p. 1224–1229.

152. Beversdorf, D.Q., et al., Timing of prenatal stressors and autism. Journal of autism and developmental disorders, 2005. 35(4): p. 471–478.

153. Courchesne, E., et al., Unusual brain growth patterns in early life in patients with autistic disorder: an MRI study. Neurology, 2001. 57(2): p. 245–254.

154. Limperopoulos, C., et al., Does cerebellar injury in premature infants contribute to the high prevalence of long-term cognitive, learning, and behavioral disability in survivors? Pediatrics, 2007. 120(3): p. 584–593.

155. Tsai, P.T., et al., Autistic-like behaviour and cerebellar dysfunction in Purkinje cell Tsc1 mutant mice. Nature, 2012. 488(7413): p. 647–651.

156. Al-Afif, S., et al., Splitting of the cerebellar vermis in juvenile rats—effects on social behavior, vocalization and motor activity. Behavioural brain research, 2013. 250: p. 293–298.

157. Bobee, S., et al., Effects of early midline cerebellar lesion on cognitive and emotional functions in the rat. Behavioural brain research, 2000. 112(1-2): p. 107–117.

158. Riva, D., et al., Gray matter reduction in the vermis and CRUS-II is associated with social and interaction deficits in low-functioning children with autistic spectrum disorders: a VBM-DARTEL Study. Cerebellum, 2013. 12(5): p. 676–85.

159. Fleming, S.M., et al., Early and progressive sensorimotor anomalies in mice overexpressing wild-type human α-synuclein. Journal of Neuroscience, 2004. 24(42): p. 9434–9440.

160. Lam, H.A., et al., Elevated tonic extracellular dopamine concentration and altered dopamine modulation of synaptic activity precede dopamine loss in the striatum of mice overexpressing human α-synuclein. Journal of neuroscience research, 2011. 89(7): p. 1091–1102.

161. Khoury, E., et al., [Sensorimotor aspects and manual dexterity in autism spectrum disorders: A literature review]. Encephale, 2020. 46(2): p. 135–145.

162. Mosconi, M.W. and J.A. Sweeney, Sensorimotor dysfunctions as primary features of autism spectrum disorders. Sci China Life Sci, 2015. 58(10): p. 1016–23.

163. Sokhadze, E.M., et al., Behavioral, Cognitive, and Motor Preparation Deficits in a Visual Cued Spatial Attention Task in Autism Spectrum Disorder. Appl Psychophysiol Biofeedback, 2016. 41(1): p. 81–92.

164. Unruh, K.E., et al., Cortical and subcortical alterations associated with precision visuomotor behavior in individuals with autism spectrum disorder. J Neurophysiol, 2019. 122(4): p. 1330–1341.

165. König, C., et al., Thirty Mouse Strain Survey of Voluntary Physical Activity and Energy Expenditure: Influence of Strain, Sex and Day-Night Variation. Front Neurosci, 2020. 14: p. 531.

166. Valagussa, G., et al., Toe Walking Assessment in Autism Spectrum Disorder Subjects: A Systematic Review. Autism Res, 2018. 11(10): p. 1404–1415.

167. Kindregan, D., L. Gallagher, and J. Gormley, Gait deviations in children with autism spectrum disorders: a review. Autism Res Treat, 2015. 2015: p. 741480.

168. Shetreat-Klein, M., S. Shinnar, and I. Rapin, Abnormalities of joint mobility and gait in children with autism spectrum disorders. Brain Dev, 2014. 36(2): p. 91–6.

169. Cook, J., From movement kinematics to social cognition: the case of autism. Philos Trans R Soc Lond B Biol Sci, 2016. 371(1693).

170. Borovac, J., M. Bosch, and K. Okamoto, Regulation of actin dynamics during structural plasticity of dendritic spines: Signaling messengers and actin-binding proteins. Mol Cell Neurosci, 2018. 91: p. 122–130.

171. Koleske, A.J., Molecular mechanisms of dendrite stability. Nat Rev Neurosci, 2013. 14(8): p. 536–50.

172. Penzes, P. and I. Rafalovich, Regulation of the actin cytoskeleton in dendritic spines. Adv Exp Med Biol, 2012. 970: p. 81–95.

173. Blanchoin, L., et al., Actin dynamics, architecture, and mechanics in cell motility. Physiol Rev, 2014. 94(1): p. 235–63.

174. Pollard, T.D. and J.A. Cooper, Actin, a central player in cell shape and movement. Science, 2009. 326(5957): p. 1208–12.

175. Yan, Z., et al., Protein phosphatase 1 modulation of neostriatal AMPA channels: regulation by DARPP-32 and spinophilin. Nat Neurosci, 1999. 2(1): p. 13–7.

176. Satoh, A., et al., Neurabin-II/spinophilin. An actin filament-binding protein with one pdz domain localized at cadherin-based cell-cell adhesion sites. J Biol Chem, 1998. 273(6): p. 3470–5.

177. Feng, J., et al., Spinophilin regulates the formation and function of dendritic spines. Proc Natl Acad Sci U S A, 2000. 97(16): p. 9287–92.

178. Arikkath, J. and L.F. Reichardt, Cadherins and catenins at synapses: roles in synaptogenesis and synaptic plasticity. Trends Neurosci, 2008. 31(9): p. 487–94.

179. Basu, R., M.R. Taylor, and M.E. Williams, The classic cadherins in synaptic specificity. Cell Adh Migr, 2015. 9(3): p. 193–201.

180. Seong, E., L. Yuan, and J. Arikkath, Cadherins and catenins in dendrite and synapse morphogenesis. Cell Adh Migr, 2015. 9(3): p. 202–13.

181. Moya, P.R., et al., Rare missense neuronal cadherin gene (CDH2) variants in specific obsessive-compulsive disorder and Tourette disorder phenotypes. Eur J Hum Genet, 2013. 21(8): p. 850–4.

182. Hussman, J.P., et al., A noise-reduction GWAS analysis implicates altered regulation of neurite outgrowth and guidance in autism. Mol Autism, 2011. 2(1): p. 1.

